# Conformational analysis of chromosome structures reveals vital role of chromosome morphology in gene function

**DOI:** 10.1101/2023.02.18.528138

**Authors:** Yuxiang Zhan, Asli Yildirim, Lorenzo Boninsegna, Frank Alber

## Abstract

The 3D conformations of chromosomes are highly variant and stochastic between single cells. Recent progress in multiplexed 3D FISH imaging, single cell Hi-C and genome structure modeling allows a closer analysis of the structural variations of chromosomes between cells to infer the functional implications of structural heterogeneity. Here, we introduce a two-step dimensionality reduction method to classify a population of single cell 3D chromosome structures, either from simulation or imaging experiment, into dominant conformational clusters with distinct chromosome morphologies. We found that almost half of all structures for each chromosome can be described by 5-10 dominant chromosome morphologies, which play a fundamental role in establishing conformational variation of chromosomes. These morphologies are conserved in different cell types, but vary in their relative proportion of structures. Chromosome morphologies are distinguished by the presence or absence of characteristic chromosome territory domains, which expose some chromosomal regions to varying nuclear environments in different morphologies, such as nuclear positions and associations to nuclear speckles, lamina, and nucleoli. These observations point to distinct functional variations for the same chromosomal region in different chromosome morphologies. We validated chromosome conformational clusters and their associated subnuclear locations with data from DNA-MERFISH imaging and single cell sci-HiC data. Our method provides an important approach to assess the variation of chromosome structures between cells and link differences in conformational states with distinct gene functions.

## Introduction

With the advent of single cell super resolution imaging^1,2^, multiplexed FISH imaging^3–6^, single cell genomics experiments^7–10^, and data driven genome modeling^11–22^ it is now possible to analyze 3D structures of chromosomes and entire genomes at single cell level. Chromatin loops, topological associated domains (TADs) and patterns of chromatin compartmentalization are readily detected in ensemble averaged Hi-C data^23–26^ but are very dynamic in nature and subsequently show large stochastic variations at single cell level^27,28^. For instance, chromatin loops, detected at specific locations in ensemble Hi-C are likely present only in 3 to 6.5% of cells at any given time^29^ and TAD domain boundaries are only rarely observed at the ensemble average position but rather stochastically distributed, because of dynamic loop extrusion processes^13,28,30,31^. Thus, detailed analysis of single cell chromosome structures are only meaningful when considering the entirety of structural variability observed in a cell population^32–34^.

Unlike chromatin loops and TADs, little research has been conducted on structural variations of long-range interactions and whole chromosome morphologies, specifically to investigate the role of these structural variations on global genome organization and gene function. Recent evidence from multiplexed FISH imaging^3,6,32^ and single cell Hi-C experiments^10,35–37^ suggests large structural variations of chromosome morphologies between single cells. These structural differences can affect spatial positions of genes within chromosome territories and thus a gene’s exposure to functional compartments and nuclear bodies, which have been shown to be of relevance for gene function^38^. For instance, transcriptional activities of individual genes can be heightened in the immediate vicinity of nuclear speckles^39–41^. However, up to this point it remains unclear if the variability in 3D chromosome morphology plays any role in the regulation and cell-to-cell heterogeneity of gene function.

In this study, we address this point by examining the cell-to-cell variability of 3D chromosome morphologies within their nuclear environment and by studying how these structural variations alter the functional microenvironment of genomic regions in the nucleus, as defined by their radial positions, or distances to nuclear speckles and lamina compartments. Because of the stochastic nature of 3D chromosome structures, several important questions emerge. First, can the structures of the same chromosome in different cells be classified into prevailing structural states that define distinct chromosome morphologies? Second, do chromosome morphologies of prevailing structural states relate to distinct functional properties of genes in these chromosomes?

To address these questions, we first introduce an approach to classify structural variations of chromosome morphologies in single cells from ensembles of 3D chromosome structures, extracted either from multiplex DNA-MERFISH imaging^3^, or structure models generated with our data-driven structure modeling approach^11^. We then study if chromosomal regions are exposed to different nuclear microenvironments in different structural states to detect potential functional variations of genes located in different chromosome morphologies.

Due to the dynamic nature of chromosome structures, their classification based on 3D coordinates is challenging, because some functionally unrelated regions can show large degree of randomness in their relative positions, overshadowing relevant structural relationships between other chromosomal regions. Our approach overcomes this problem and transforms the problem of classifying individual chromosomes structures to a problem of detecting maxima of a density distribution in a reduced 2-dimensional space, where each data point represents a chromosome conformation and the detection of local maxima in the probability density function determines locations of highly occupied clusters of chromosomes with similar 3D conformations. Thus, our approach determines subpopulations of chromosomes with similar 3D morphology. We discovered that a given chromosome can be clustered into around 5 to 10 morphology classes, which are distinguished by the presence or absence of characteristic chromosome territory domains that vary in their relative locations to each other. The boundaries of these territory domains play a fundamental role in establishing conformational variation and their sequence locations are shared across various cell types (GM12878, H1-hESC and HFFc6). We validated the observed chromosome conformational states and chromosome territory domain boundaries with data from multiplex DNA-MERFISH imaging data^3^ and single cell sci-HiC experiments^10^. We then discovered that distinct chromosome morphologies (i.e. conformational states) favor certain nuclear locations of some chromosomal regions and thus modulate the functional properties of these genomic regions. For instance, preferences in radial positions, distances to the nearest speckle, nucleolus and lamina differ substantially between the same chromosomal regions in different conformational states. These observations point to functional differences of chromatin in different conformational states, as smaller distances to nuclear speckles are typically associated with increased transcriptional activities of genes. Our observations therefore indicate that chromosome morphologies can play a key role in modulating functional properties of some chromosomal regions, and can, at least partially, be responsible for the cell-to-cell heterogeneity of the expression of some genes. Our method provides an important approach to study chromosome conformational variations and reveal links between conformational states of chromosomal regions and gene functions.

## Results

### Structure generation

We first apply our approach for characterizing chromosome morphologies and their functional qualities to an ensemble of diploid 3D genome structures that were generated at 200kb resolution from Hi-C data^11,13,25,26^. Our population-based 3D genome modeling method (Integrative Genome Modeling, Methods)^11,13,42,43^ provides us with a large sampling of 10,000 diploid 3D genome structures per cell type, which, as a whole, reproduce Hi-C data and predict with high accuracy other orthogonal experimental data^11,13^, namely average radial positions of genomic regions from GPseq experiments^44^, mean speckle distances from SON TSA-seq^45^, mean distances and contact frequencies to the nuclear periphery from lamin B1 TSA-seq^45^ and lamin B1 pA-DamID^46^, respectively. Moreover, predicted chromosome structures show good agreement with single cell 3D chromosome conformations from multiplex DNA-MERFISH experiments^3,11,13^, and also reproduce with good accuracy speckle and lamina association frequencies of genomic regions from DNA MERFISH^11,13^. We first focus our analysis on genome structure models from lymphoblastoid cells (GM12878) from previously published work^13^, human fibroblast (HFFc6)^11^ and human embryonic stem cells (H1ESC)), before classifying genome structures from DNA-MERFISH experiments^3^.

### Approach

To classify chromosomes based on their morphology, we first extract individual chromosomes of a given type from each of the whole genome structures in the cell population. Both homologous chromosome copies in the diploid genome are selected, resulting in a total of 20,000 chromosome structures for each autosome (**Fig. 1**). For each 3D chromosome structure, we then construct a distance matrix, which then serves as input into our dimension reduction and clustering scheme (**Fig. 1**). We then use a two-step dimension reduction approach to cluster chromosome structures based on their distance matrices into conformational states (Methods). Specifically, each normalized distance matrix is represented as a 2D image. Our two-step process then combines a convolutional autoencoder (consisting of an encoder and a decoder module) with a dimension reduction step using t-distributed stochastic neighbor embedding (t-SNE)^47^ (Methods). The encoder module reduces a distance matrix to a latent vector that can reconstruct the original matrix by the decoder module. The method reduces dimensions, while preserving enough information to reconstruct the original image. To construct a convolutional autoencoder we use convolutional layers, max pooling layers and up sampling layers, which is frequently used for image embedding and classification (**Supplementary Fig. 1**) (Methods). t-SNE, a method to separate data points in a reduced data space, is then used to map the latent vectors (generated by the autoencoder) to a lower dimensional space (**Fig. 1**). Finally, we use a kernel density estimation to calculate a probability density function (pdf) that represents the chromosome conformational space in the t-SNE reduced dimensions. The resulting density probability matrix shows a balanced distribution of local maxima separated by deep valleys (**Fig. 1** lower panels), indicating the presence of a number of preferred conformational states per chromosome, which are subsequently identified by a segmentation algorithm (Methods).

**Figure 1:**
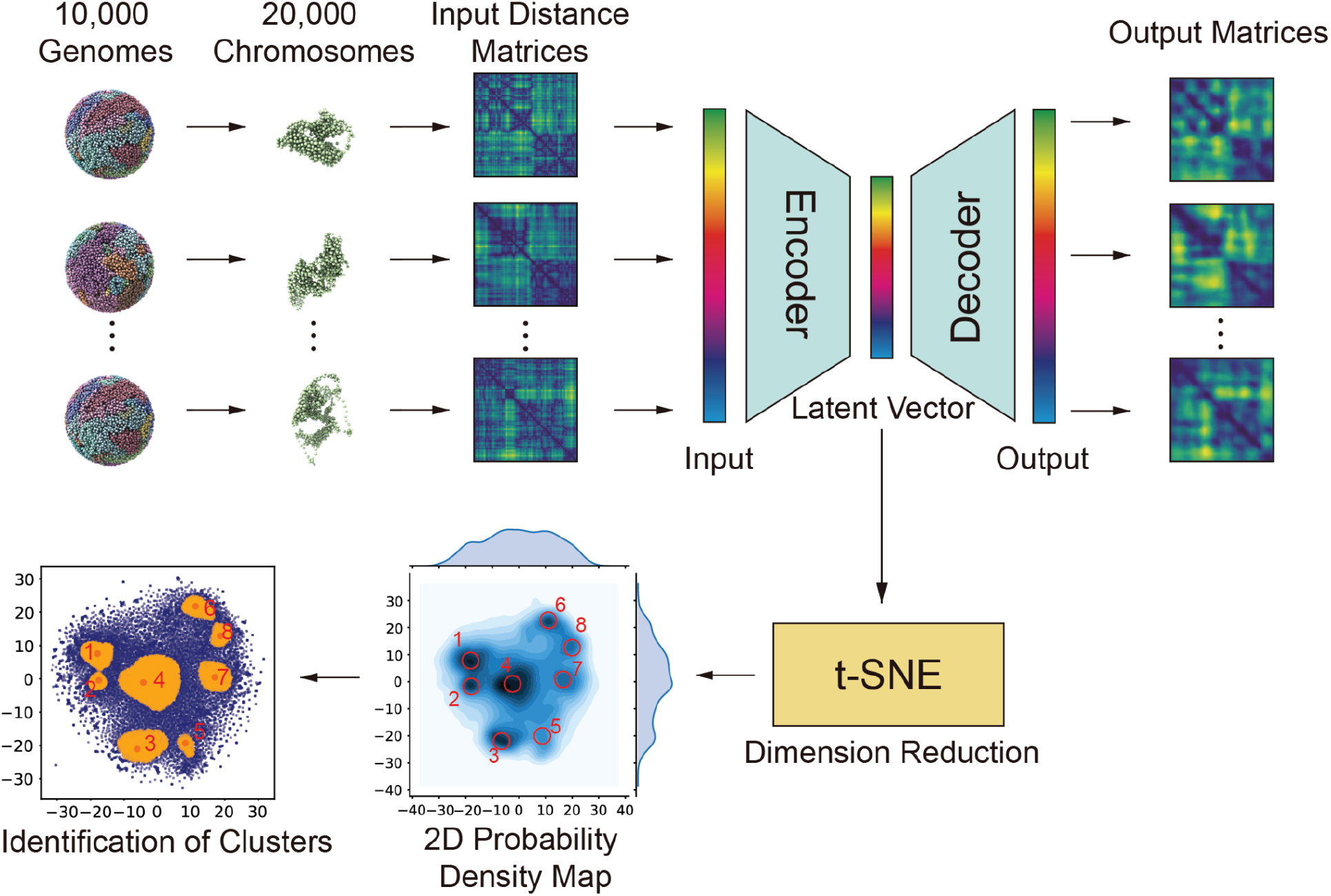
Overview of the two-step dimension reduction. Every chromosome structure is represented by an input distance matrix, which is constructed by calculating pairwise Euclidean distances between each pair of loci in the chromosome structure. After preprocessing, the matrix is then used as the input of the autoencoder. After minimizing the loss between input matrices and output matrices, the latent vectors are then embedded by t-SNE^47^ to obtain a distribution of all chromosome structures in 2D space. The resulting distribution is further used for peak detection and identification of clusters of chromosome structures (Methods).

We also tested other clustering methods. However, principal component analysis (PCA), multidimensional scaling (MDS)^48^, locally linear embedding (LLE)^49^, isomap^50^ and spectral embedding (SE)^51^ methods are unable to determine distinct clusters with chromosomes of similar conformational morphology, while uniform manifold approximation & projection (UMAP)^52^ and T-distributed stochastic neighbor embedding (t-SNE) (applied directly to distance matrices alone) produced unbalanced clusters, in which the majority of structures were part of only a single cluster (**Supplementary Fig. 2**). Instead, the balanced distribution of local maxima in the resulting density probability matrix of our two-step clustering approach (**Fig. 1** lower panels) indicates the presence of a number of preferred conformational states per chromosome.

### A large fraction of chromosome structures can be clustered into a few conformational states

We then identify clusters of similar chromosome conformations by determining local maxima in the probability density distribution as cluster centers and identify structures associated to each cluster center by watershed segmentation of the probability density distribution (**Fig. 1** lower left panel, and Methods). Chromosome structures part of the same segmentation are in the same conformational cluster. For chromosome 6 about 40% of all chromosome structures can be clustered into 8 dominant conformational clusters (**Fig. 1**, lower left panel, **Fig. 2A**). The occupancy of each cluster is defined by the number of structures in a cluster divided by the total number of all clustered chromosome conformations. The occupancy among clusters varies (**Fig. 2A**). For chromosome 6, cluster 4 has the highest occupancy containing ~40% of all the clustered structures, while all other clusters each occupy less than 20% of all clustered structures. Similar results are found also for other chromosomes (**Supplementary Fig. 4**).

**Figure 2:**
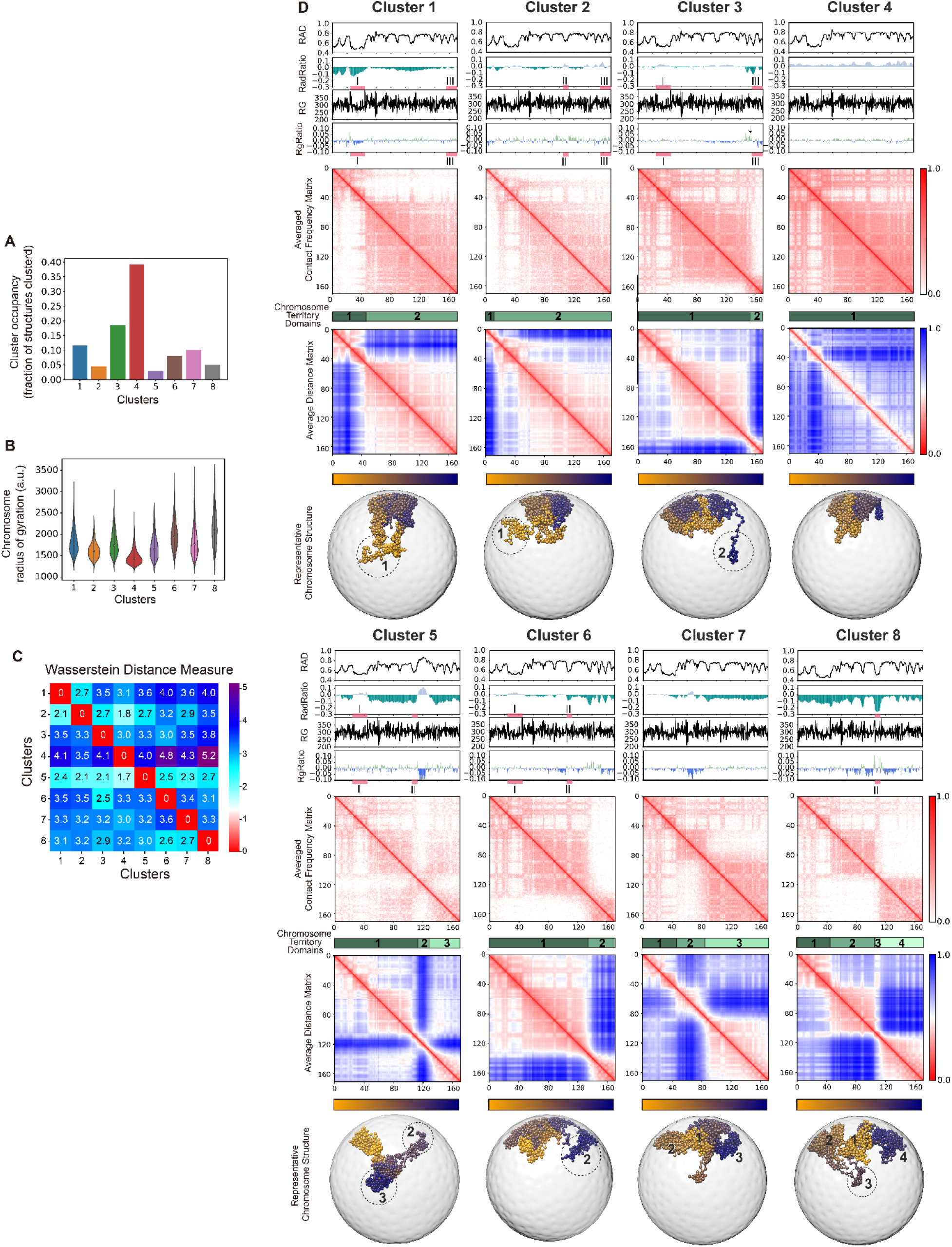
Clustering of chromosome 6 structures reveals dominant chromosome morphologies. **A**, The cluster occupancy of the 8 predicted clusters for chromosome 6. The occupancy is defined by the fraction of structures in each cluster. **B**, The distributions of the radius of gyration of all structures in each cluster (Methods). **C**, Pairwise dissimilarity measure between chromosome structures in 8 clusters. The dissimilarity matrix is calculated by measuring the average Wasserstein distance^53^ of all intra-chromosomal distance distributions between two clusters. Each entry represents the log fold ratio between the inter-cluster dissimilarity and the intra-cluster dissimilarity, where positive values indicate the inter-cluster dissimilarity is larger than the intra-cluster dissimilarity. **D**, For each cluster the following information is shown (Methods): (top panel) The average radial position profile (RAD) calculated from all all chromosome structures in the cluster; (second panel from top) RadRatio: the log fold ratio of the average radial position in the cluster with respect to full ensemble average; (third panel from top) RG: the radius of gyration of a 1Mb chromosomal region centered at the target locus (fourth panel from top) RGRatio: the log fold ratio of RG in the cluster with respect to the value in the full ensemble; (fifth panel from top) The average contact frequency matrix calculated from all structures in a cluster; (sixth panel from top) different shade of green indicate the location of chromosome territory domains; (seventh panel from top) The average distance matrix calculated from all chromosome structures in the cluster; (eighth panel from top) Selected example of a chromosome structure in the cluster. Numbers and circles indicate chromosomal regions of the corresponding chromosome territory domains. The color bar indicates the sequence position of each chromosomal region. We also highlighted several specific genomic regions (regions I, II and II) below the RadRatio and RGRatio profiles, which are compared and discussed in the text.

### Preferred conformational states define distinct chromosome morphologies

While chromosomes within each cluster share similar morphology, chromosomes between clusters differ in their structures. For instance, when we measure the compactness of a chromosome structure by calculating its radius of gyration, it is apparent that the structures of each cluster vary largely in their shapes (**Fig. 2B**). Chromosomes in cluster 8 show the lowest compaction with an average radius of gyration that is about 50% larger than chromosomes in cluster 4, which show the most compact structures (**Fig. 2B**). Overall, chromosomes in clusters with relatively low compaction, and thus, highest radius of gyration show also the largest variations of their compaction values within the cluster (e.g., clusters 6, 8). Moreover, cluster 4 containing chromosomes with the most compact structures shows the highest occupancy (**Fig. 2AB**).

Next, we quantify the structural similarity between chromosome structures within and between clusters by calculating the Wasserstein distance^53^ between their pairwise chromatin distance distributions. Specifically, we measure the difference between distributions of all intra-chromosomal distances calculated from all chromosomes in each cluster. In other words, for a given pair of chromosomal regions *i* and *j*, we calculate the distance distribution from all chromosomes in a cluster and assess its similarity with the corresponding distance distribution calculated from the structures in another cluster. The similarity between two such distance distributions is calculated by their Wasserstein distance metric^53^. The combined distance measure between two clusters is then defined as the average of all Wasserstein distances between all intra-chromosomal distance distributions calculated from the chromosomes in the two clusters (**Fig. 2C**). We normalize this measure by the average Wasserstein distance for chromosome structures within the same cluster (Methods). We observe that the average Wasserstein distance is always substantially larger (i.e., ~2-4 fold) for structures in different clusters, showcasing the structural distinction between chromosomes in the different clusters. We also found similar results when assessing clusters with other distance measures, including a Euclidean distance measure and Gaussian dissimilarity^32,54^, confirming an overall higher similarity between structures within than between different clusters (Methods) (**Supplementary Fig. 3**).

Importantly, each cluster shows distinct average contact frequency matrices, calculated from the physical chromatin contacts of all chromosomes in each cluster (**Fig. 2D**, average contact frequency matrix shown in fifth panel from top). These distinct contact patterns confirm the presence of characteristic chromosome morphologies in each cluster. A characteristic feature of these contact patterns is the presence of domain boundaries that divide the chromosome territory into large units, with increased interaction frequencies within, and reduced contact frequencies between territory domains (for instance in clusters 1, 3, 6 and 8 of chromosome 6, **Fig. 2D** (territory domain locations are indicated by green blocks below the contact frequency matrix.)) These territory domains are particularly evident when calculating the average distance matrix for each cluster (i.e., from all intra-chromosomal 3D distances in a chromosome), because territory domains show increased spatial distances from each other, thus are separated spatially from each other in 3D space (see for instance domains 2 in clusters 3 and 6 in **Fig. 2D**). The spatial separation between domains explains the reduced contact frequencies between the territory domain regions. The location and size of territory domains vary between the clusters. On average, chromosome structures contain between 1 and 4 territory domains per studied chromosome (green blocks below in **Fig. 2D, Supplementary Fig. 4DH**). **Figure 2D** also shows representative structures of the chromosome morphology found in each cluster of chromosome 6. The structures show the spatial separation between territory domains, as observed in the average distance matrices. For instance, cluster 6 of chromosome 6 shows a territory domain boundary at around 134 Mb sequence position, which separates the q-arm terminal end of the chromosome into a separate territory domain (domain 2 in cluster 6 (134-171 Mb), **Fig. 2D**). This domain shows relatively increased spatial distances and low chromatin contact frequencies to other chromosomal regions upstream of the territory domain boundary (**Fig. 2D**). In contrast, cluster 3 shows a domain boundary at sequence position 155 Mb, which forms an even smaller domain at the q-terminal end of chromosome 6 (domain 2 of cluster 3 (155-171 Mb), **Fig. 2D**), which is well separated from the bulk of the remaining chromosome (also evident in the representative chromosome structure (lower panel).) Cluster 2 contains a relatively small chromosome territory domain at the p-arm terminal end of chromosome 6 (domain 1 in cluster 2 (0-10 Mb), **Fig. 2D**). Noticeable, the boundaries between territory domains act as hinge regions allowing the relative positions of territory domains to vary in 3D between models, while the territory domain itself appears as structural units (representative structures in **Fig. 2D**). Accordingly, chromosomes in different clusters also show differences in their local chromatin compaction, as measured by the radius of gyration (RG) over a 1 MB window of the chromatin fiber (**Fig. 2D**, third profile panel from top). Local peaks in the RG profile are regions with relatively low fiber compactness (which often correspond to TAD boundaries as previously shown^11,13^). These profiles show distinct differences between clusters, noticeable at locations of some territory domain boundaries. To highlight these differences, we calculated the RGRatio as the log ratio of the average RG value for a chromatin region in the cluster and the overall ensemble. For instance, the RGRatio profile of both domain 3 boundaries in cluster 8 show high values, indicating that these boundary regions are decompacted in cluster 8 in comparison to the same region in the overall ensemble (**Fig. 2D**, RG and RGRatio profiles are shown in the third and fourth panel from top.) Also, the boundary that separates the small domain 2 in cluster 3 from the bulk of the chromosome territory shows substantially increased RGRatio, thus shows a substantial decrease in fiber compactness in the cluster in comparison to the compactness in the overall ensemble of chromosomes, therefore allowing domain 2 the freedom to loop away from the bulk of the remaining chromosome territory (see black arrow in RGRatio of cluster 3 in **Fig. 2D**). These observations indicate that local chromatin properties can facilitate the formation of specific chromosome morphologies.

Besides chromosome 6, also other chromosomes show very similar results, with distinct chromosome contact frequency patterns for each cluster. For instance, conformational clusters for chromosome 8 and chromosome 10 are also distinguished by a total of 8-10 different chromosome morphologies with distinct territory domains whose locations vary between individual clusters (**Supplementary Fig. 4**).

### Chromosome clusters can be validated by imaging experiments

We assessed our findings with data from multiplex DNA-MERFISH imaging experiments^3^, which traced 3D coordinates of whole diploid genomes in IMR90 fibroblast cells at a step size of ~3Mb. To allow a direct comparison, we down sampled our models to the genomic regions sampled in the experiment and classified the chromosome conformations from DNA MERFISH into clusters (Methods), based on the similarity of their distance matrix with cluster averages in our models (**Fig. 3AB**). For chromosome 6, around 60% of all chromosome structures from DNA MERFISH (about 4,000 structures) can be classified into our predicted chromosome clusters based on their structural similarity (**Fig. 3A-E)**. When we average the distance matrices of all imaged chromosome structures in each cluster, we see that the cluster averages from DNA-MERFISH experiments are almost identical to those calculated from our models (**Fig. 3AB**). Also in experiment, cluster 4, with the most compact chromosome structures, shows the highest occupancy (**Fig. 3F**). Moreover, individual representative single cell chromosome structures from DNA-MERFISH imaging show almost identical chromosome structures and distance matrices to those from our predicted models (**Fig. 3CD**). Thus, DNA-MERFISH imaging confirms the presence of preferred chromosome morphologies and the presence of chromosome territory domains that vary in their locations between the clusters.

**Figure 3:**
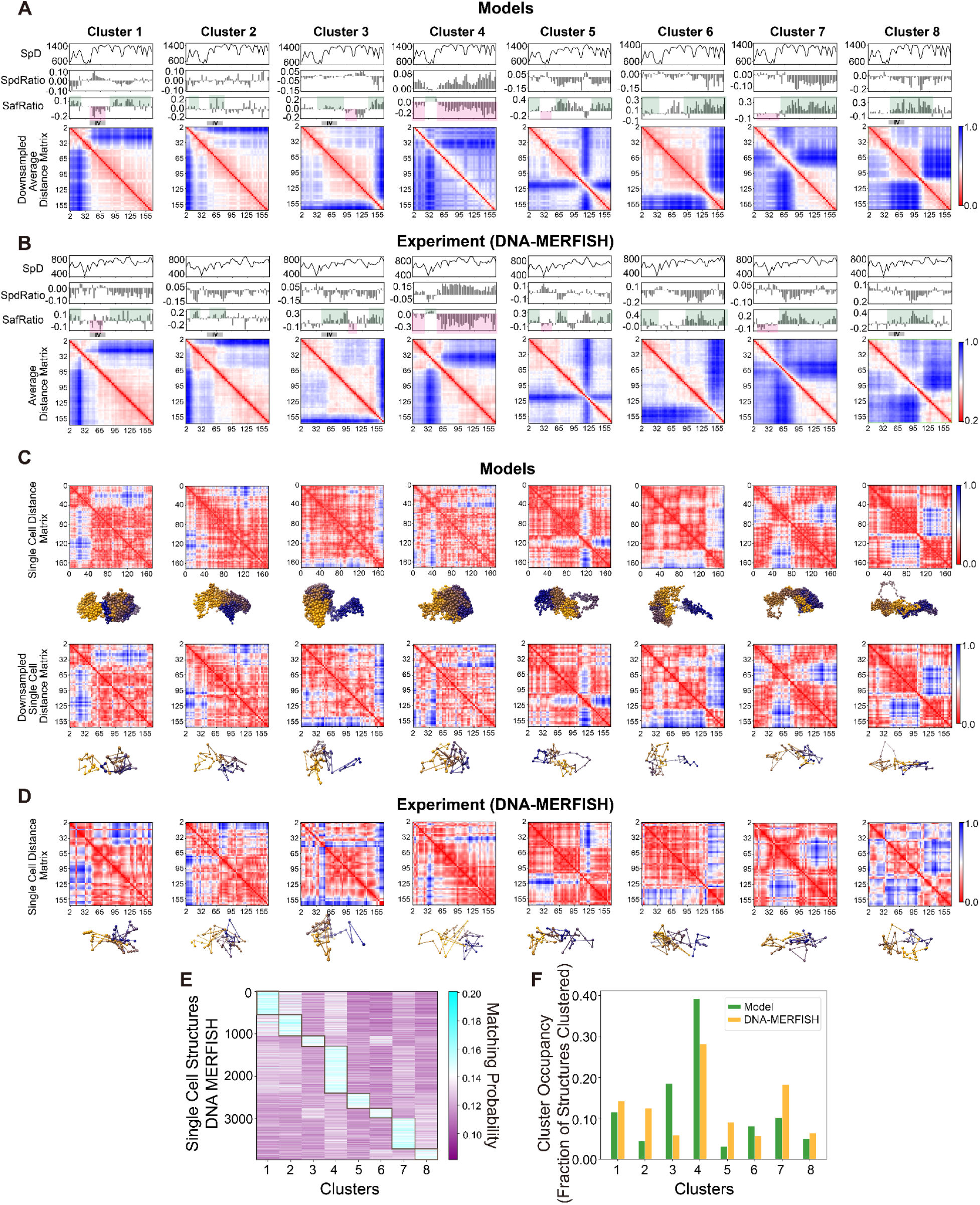
Chromosome clusters can be validated by imaging experiments. **A**, Average distance matrices from modeled chromosomes (chromosome 6) in each cluster downsampled to the respective coverage as observed in DNA-MERFISH experiments^3^. Tick labels of all distance matrices indicate sequence location in Mb. Also shown are several structural features calculated from the genome structures of the cluster ensemble, namely the average distance to the nearest speckle (SpD), the log fold ratio of the average distance to nearest speckle (SpdRatio) in the cluster with respect to full ensemble average and the log fold ratio of the speckle association frequency (SafRatio) in the cluster with respect to the full ensemble average. Green and magenta shaded areas indicate coarse matches of the varied SafRatio in predicted and experimental determined clusters. For comparison with experiment distance matrices were down sampled to the same coverage in the experiment (window size 3Mb). **B**, The corresponding average distance matrices of chromosomes from DNA-MERFISH experiments for each cluster. Shown are also the average distance to the nearest speckle (SpD) in each cluster, the log fold ratio of the average distance to nearest speckle (SpdRatio) in each cluster and the log fold ratio of the speckle association frequency (SafRatio) in each cluster, all measured from DNA-MERFISH imaging^3^. Green and magenta shaded areas indicate coarse matches of the varied SafRatio in predicted and experimental determined clusters. **C**, Selected representative examples of single cell modeled structures and the corresponding downsampled version for chromosome structures for each cluster. Matrices are calculated with window size 200kb. **D**, Average distance matrices and selected representative single cell chromosome structures from DNA-MERFISH experiment for each cluster. In panels A, B, lower panel C and D distance matrices are shown at 3Mb resolution (coverage in DNA MERFISH experiment); in upper panel C the distance matrix is shown at 200 kb resolution. **E**, Matching probabilities indicating the similarities between distance matrices of all classified single cell DNA-MERFISH chromosome structures against all modeled clusters. Note that around 60% of the single cell structures are successfully classified and assigned to one of the modeled clusters. **F**, Comparison of the cluster occupancy between the chromosome conformational clusters observed in our models and corresponding chromosome conformational clusters from DNA-MERFISH experiments.

### Chromosome morphologies show distinct preferences in nuclear locations of chromosomal regions

We now focus on the nuclear organization of chromosomes in different morphologies. The question we want to address is: does the morphology of chromosome structures relate to specific nuclear locations of chromosomal regions and thus modulate their functional properties? Because we model whole genome structures, we can analyze chromosome structures in their nuclear context. First, we can extract the nuclear radial positions of chromosomal regions and average the radial positions from chromosomes in each conformational cluster (**Fig. 2D**, top profile panel). We assess the differences of the radial (RAD) profile between clusters by calculating the RadRatio, defined as the log ratio between the average radial position of a genomic region (RAD) in a cluster and its value in the whole population of clustered structures (**Fig. 2D**, second profile panel from top). A negative RadRatio value for a genomic region indicates that its average radial position in the cluster is closer to the nuclear interior in the cluster than in the overall population as a whole. We see that RadRatio profiles differ substantially between clusters for chromosome 6 in particular for clusters 1, 3, 5, 6 and 8, which show pronounced peaks in the RadRatio profile, both in positive and negative values. For instance, the RadRatio profiles in clusters 1 and 3 differ substantially across the entire chromosome (**Fig. 2D**). In cluster 1 the genomic location 24-48 Mb (region **I** of cluster 1 in **Fig. 2D**), which includes the MHC gene cluster, is substantially shifted towards interior nuclear positions (i.e. negative RadRatio values) in comparison to the location of the same region **I** in clusters 3, 5 and 6, which show positive RadRatio values, thus more exterior locations than in the population average (p-values of Welch’s t-test^55^ on average radial position against clusters 3, 5 and 6 = 4.06e-12, 1.02e-03 and 6.17e-09 (Table 1), **Fig. 2D**). Instead, cluster 3 shows a small chromosome territory domain at the q-terminal chromosome end, readily visible in the contact frequency and distance maps (genomic location 155-171 Mb: domain 2/region **III** in cluster 3 in **Fig. 2D**). This territory domain loops towards the nuclear interior in cluster 3, as shown in the RadRatio profile and the representative structures (**Fig. 2D**). In other clusters (e.g., cluster 1 and 2) the same region **III** shows a more exterior nuclear location and does not form a separate domain, but is part of an overall larger chromosome territory domain (see region **III** in clusters 1 and 2, **Fig. 2D**). Another example is the genomic region **II** at sequence location 105-114 Mb. It forms a small territory domain in cluster 8 (domain 3, region **II** in cluster 8 in **Fig. 2D**), which shows strongly negative RadRatio, and therefore loops towards the nuclear interior. Instead, the same region **II** in cluster 6 is part of a larger territory domain, which restricts its looping towards the nuclear interior in comparison to cluster 8 (p-values = 1.04e-09, Welch’s t-test on average radial positions in cluster 8 and 6 (Table 2)). In cluster 2 the same region **II** is even shifted more towards the nuclear periphery in comparison to the population average (see negative RadRatio for regions **II** in cluster 2 in **Fig. 2D**)(p-values = 4.95e-17, Welch’s t-test on average radial positions in cluster 8 and 2 (Table 2)). Moreover, in cluster 4 chromosomes are in their most compact conformation, which correlates with a shift of almost the entire chromosomal regions towards the nuclear periphery in comparison to the population average (see positive values in RadRatio profile in **Fig. 2D**) (p-values = 1.69e-34, Welch’s t-test on average radial positions in cluster 8 and 4 (Table 2)).

**Table 1:**
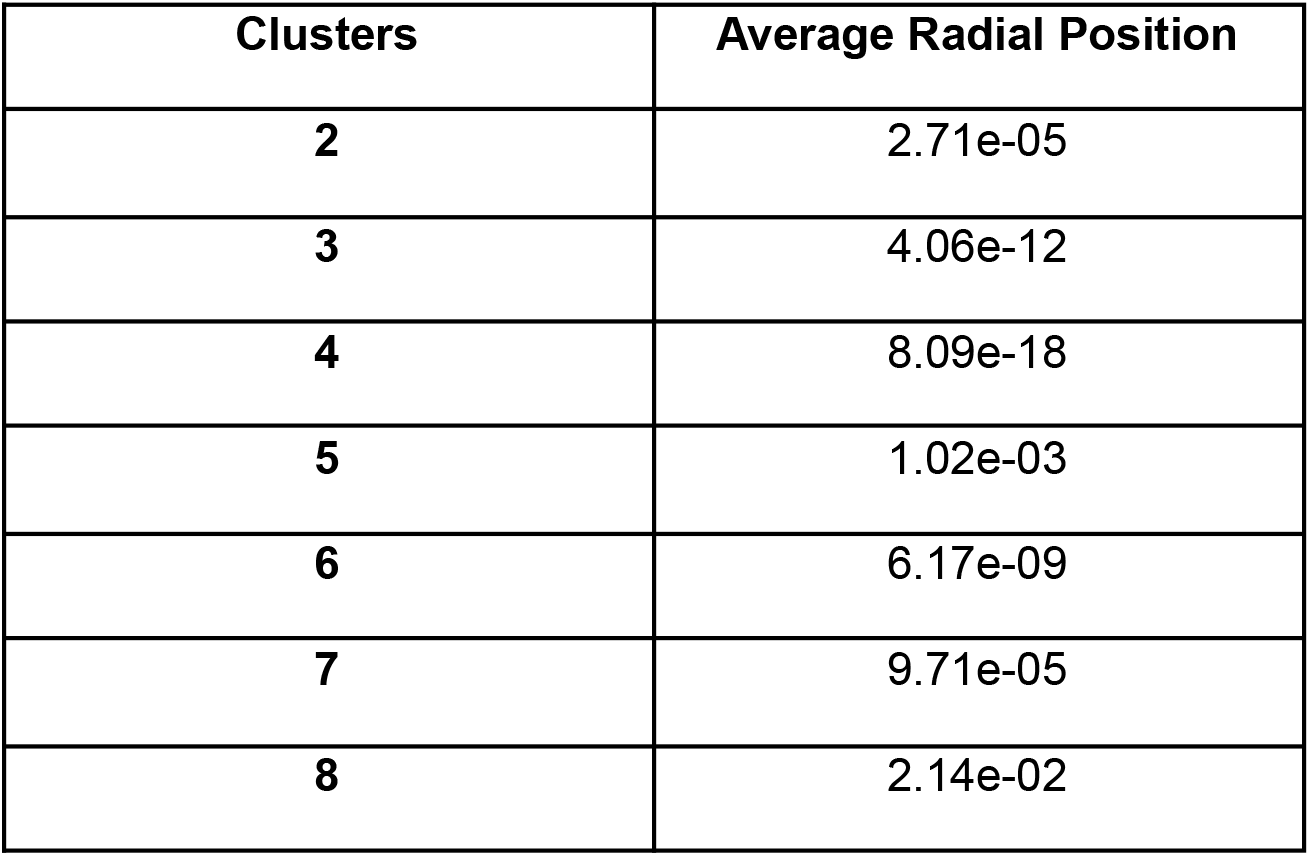
P-values of the two-sample t-test (Welch’s t-test)^55^ between cluster 1 and the other clusters of Chr6 on average radial position of region I (24-48 Mb)

**Table 2:**
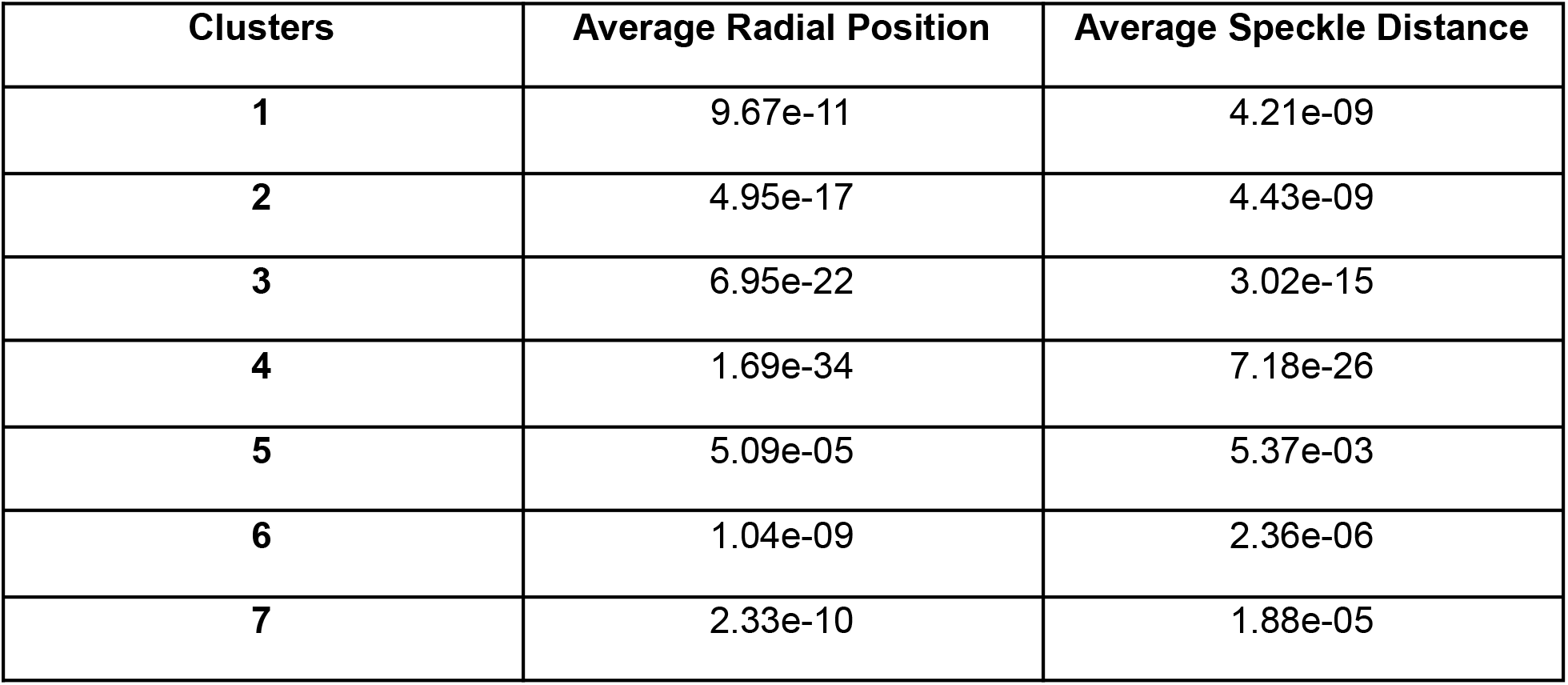
P-values of the two-sample t-test (Welch’s t-test)^55^ between cluster 8 and the other clusters of Chr6 on average radial position and average speckle distance of region II (105-114 Mb)

### Chromosome morphologies are likely linked to differences in gene functions

The differences in radial positions of genomic regions could indicate differences in their functional properties. It is known that transcriptionally active chromatin is more likely located towards the nuclear interior, and highly transcribed genes are often found close to nuclear speckles^3,23,56^. It has been previously shown that smaller mean speckle distances of genes correlate with high transcriptional activity^13,39–41^. We therefore examine if chromosome conformations influence speckle associations of genomic regions. We previously showed that our models can predict with good accuracy locations of nuclear speckles in single cell models^11,13^ by identifying the geometric centers of highly connected clusters of speckle-associated chromatin in each model. Our models predicted with good accuracy data from SON TSA-seq experiments^45^, which measure the mean distances of genomic regions to speckles (0.87 Pearson’s correlation coefficient between predicted and experimental data^13^), as well as speckle association frequencies (SAF) of genomic regions from DNA-MERFISH imaging experiments (0.79 Pearson’s correlation coefficient between prediction and experiment^13^). When we predicted SAF, SON-TSA-seq and mean speckle distances (SpD) for chromosomes in different clusters, we noticed that all these profiles vary considerably for chromosomes in different clusters (**Fig. 4A**, top 3 panels). For instance, in cluster 8 the more interior nuclear location of chromosomal region **II** (genomic location: 105-114 Mb) (**Fig. 4A**) leads to a substantially decreased mean speckle distance (**Fig. 4B)**, as shown by the higher predicted SON TSA-seq signals and a ~four times larger SAF value in comparison to the same region in cluster 6 (**Fig. 4A)**, which is buried in a larger chromosome territory (p-value = 2.36e-06, Welch’s t-test on average speckle distances between cluster 6 and 8 (**Fig. 4B**, Table 2). We showed previously that a higher SAF, thus smaller mean speckle-distance, generally correlates with higher transcriptional activity^13^). We therefore speculate that in cluster 8 genes in this region (105-114 Mb – region **II** in **Fig. 4A**), if transcriptionally active, are predisposed for higher transcriptional activity than the same regions in cluster 6 (**Fig. 4A**). Moreover, chromosomes in cluster 4 have the most compact structures and almost all chromosomal regions, with the exception of the MHC gene cluster, are predicted to have larger speckle distances and decrease speckle association frequencies (SAF) in comparison to chromosomes in other clusters (**Fig. 3A**). We validated our predicted speckle distances and SAF values using the clustered chromosome conformations taken from DNA-MERFISH imaging^3^, which also image speckle locations. These images confirm that the mean speckle distances and SAF profiles vary considerably for chromosomes in different clusters (**Fig. 3B**). For a better comparison, we calculated the SafRatio, defined as the log ratio between the SAF of a genomic region in the cluster and the overall value in the ensemble of all clustered structures. The most compact conformations in cluster 4 shows indeed a more exterior nuclear location and reduced speckle association frequencies for regions across the entire chromosome, except the MHC genes, in comparison to more extended chromosome conformations in all other clusters, confirming our predictions (see reduced SafRatio (Methods), **Fig. 3B**). For instance, cluster 1 shows a distinctly different SafRAtio in comparison to cluster 4 and 7, with certain chromosomal regions showing opposing SafRatio values, indicating that these regions show different speckle distances in different clusters. For instance the p-terminal end of chromosome 6 in cluster 1 (0-24 Mb (indices 0-8) in **Fig. 3AB**) shows substantially increased SAF over the population average in both the predicted and experimental cluster 1, while the same region in cluster 4 and 7 show decreased speckle association frequency. This observation may indicate an increased transcriptional propensity for genes in the region (0-24 Mb (indices 0-8)) in cluster 1. Moreover, region IV (indices 14-20) in cluster 1 shows decreased speckle association frequencies in both experiment and prediction (SafRatio in **Fig. 3AB**), while the same region shows increased speckle association frequency in clusters 2, 3 and 8, in both the experiment and the prediction. Overall, the experimental structures confirm the predicted patterns of increased and reduced speckle associations across the chromosome clusters (see magenta and green shaded blocks in SafRatio of **Fig. 3AB**), even though the experiments were done on IMR90 cells with a different nuclear shape. In summary, our analysis reveals that changes in chromosome conformations can be linked to changes in specific nuclear locations of genomic regions relative to nuclear bodies, which possibly affect their functional properties.

**Figure 4:**
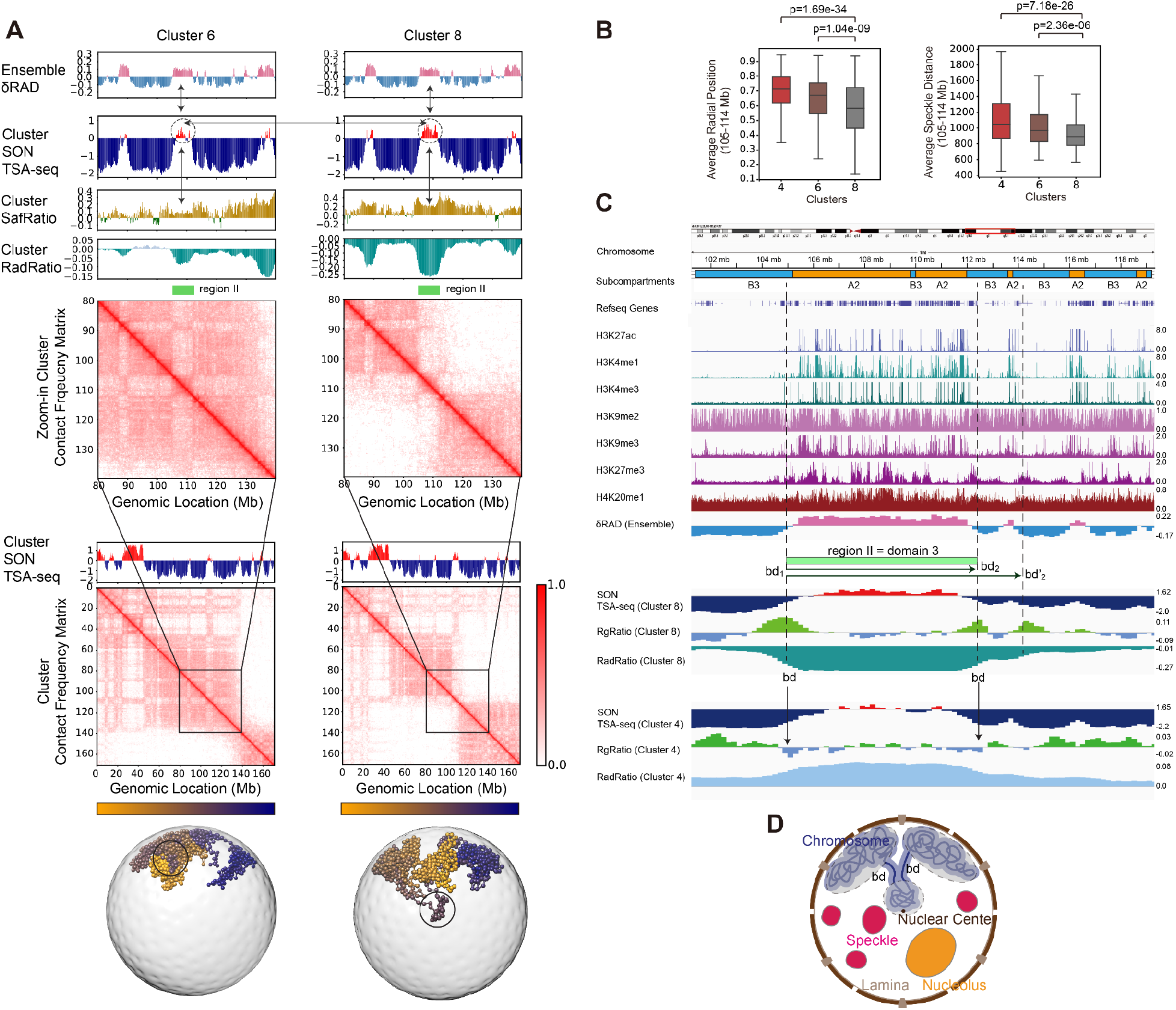
Chromosome morphologies show preferences in nuclear locations. **A**, Contact frequency matrices and 4 profiles of different structural properties for chromosome 6 in clusters 6 and 8. (Top panel) The structural variability (δRAD) of chromosomal regions (Methods). Positive values in red indicate regions of high structural variability in the ensemble of all structures. Negative regions in blue indicate regions with low structural variability in the ensemble. (Second panel from top) SON TSA-seq predicted from genome structures in each cluster. Positive values indicate shorted mean distances to nuclear speckles. (Third panel from top) SafRatio, log ratio of SAF calculated from chromosomes in the cluster over SAF calculated from structures in the whole ensemble (Methods). (Fourth panel from top) RadRatiocalculated from chromosomes in the cluster, RadRatio is defined as in Figure 2 and Methods. (Fifth panel from top) Average contact frequency matrices calculated from structures in the cluster. The first five panels show data for a zoomed-in genomic region in chromosome 6. The sixth and seventh panels from top show the predicted cluster SON TSA-seq and cluster average contact frequency matrices for the full length chromosome 6. The bottom panel shows selected representative structures for chromosome 6 in each cluster. Also indicated is region II (genomic location (105-114 Mb)), which is discussed in the text. **B**, Distributions of the average radial position and average speckle distances for a region II (genomic location: 105-114 Mb) in clusters 4, 6 and 8 as well as the p-values of Welch’s t-test^55^ between two pairs of clusters. Differences between clusters of the radial positions and average speckle distances are significant. **C**, Characteristic features for a chromosomal region spanning across region II, which forms territory domain 3 in cluster 8 and the same region II in cluster 4. Shown are epigenetic marks and other features for this sequence region. From the top to the bottom, the displayed features are chromosome sequence location, Hi-C subcompartments, refseq genes, H3K27ac, H3K4me1, H3K4me3, H3K9me2, H3K9me3, H3K27me3, H4K20me1, and the ensemble structural variability (δRAD) calculated from all structural models. In addition the following features are shown for the same regions calculated from cluster 8 and cluster 4: SON TSA-seq, RgRatio and RadRatio (Definitions as in Figure 2 and 4, Methods). Also shown are lines that indicate the territory domain in cluster 8 and corresponding domain boundaries that overlap with regions of reduced chromatin compaction (RGRatio) (bd1 and bd2). For bd2 two alternative boundaries exist in the cluster (bd2 and bd2’). **D**, Illustration of schematic features of a chromosome morphology with three territory domains. Shown are also nuclear bodies. bd regions indicate domain boundaries that show increased decompaction of the chromatin fiber (i.e., RG) in comparison to the ensemble average, and which allow the territory domain to loop towards the nuclear interior, while other territory domains remain at the periphery.

### Characteristic features of chromosome territory domain boundaries

Next, we investigate the characteristic properties that define a territory domain (TD) border (**Fig. 2D**). We noticed that if a territory domain undergoes a major change in nuclear position in a cluster (i.e., RadRatio shows pronounced minima or maxima) its boundaries show lower chromatin compaction to facilitate the passage of the domain to the interior (or exterior) regions of the nucleus (**Fig. 4AC**). The local compaction of the chromatin fiber can be calculated from the radius of gyration (RG) of a chromosomal region (Methods): The log ratio between the RG profile in a cluster over the population average (RgRatio) shows a sharp peak when a TD boundary is present in a cluster, as is shown for cluster 8 (**Fig. 4AC**) (Methods). Thus, these boundary (bd) regions show substantially reduced chromatin compaction in chromosomes of cluster 8 than in the population average (**Fig. 4C**). Interestingly, boundaries of domain 3 (region II) in cluster 8 seem to have an alternative second downstream boundary (bd_2_’), which allows the inclusion of a small gene cluster into the domain in some structures (**Fig. 4C**). In contrast, in cluster 4, where there are no domain boundaries present at these positions, chromosomes do not show any decompaction at the corresponding chromosomal regions, on the contrary, they are slightly more compacted with lower RG values than observed in the population average (RgRatio < 0, RGRatio is defined as the log ratio of the RG value for a genomic region in the cluster and its value in the ensemble (Methods) (**Fig. 4C**).

Interestingly, TD boundaries are often located close to transitions between gene poor and gene rich chromosomal regions (**Fig. 4C**). The territory domain itself contains chromatin with histone modifications related to active chromatin (e.g. H3K27ac, H3K4me1 and H3K4me3) (**Fig. 4C, Supplementary Fig. 5B, 6B, 7B**). These domains have positive SON TSA-seq signals of intermediate strength, indicating that these regions can be close to nuclear speckles in some structures. The domain boundary contains genes lacking H3K27ac and other activating histone modifications but instead are marked often with repressive histone modifications (such as H3K27me3), which mark the boundary to gene poor regions outside the domain (**Fig. 4C**). Also, domain boundaries often mark transitions between Hi-C subcompartments, as defined by Rao et al^25^. For instance, most territory domain boundaries in chromosome 6 coincide with boundaries between the A2 and B3 subcompartments. Finally, we also noticed that territory domains are also chromosomal regions with generally high cell-to-cell variability in their radial positions (see δRAD profiles in **Fig. 4C**).

### Inter-chromosomal interactions are specific to chromosome clusters

Chromosome morphologies influence the predisposition of chromosomal regions to form inter-chromosomal interactions. In some morphologies chromosomal regions may be shielded from inter-chromosomal interactions, while the same region in another morphology may be exposed to other chromosomes. To quantify the role of chromosome morphology on inter-chromosomal interactions, we calculated for each cluster the inter-chromosomal proximity matrix, as the frequency of a genomic region to be in spatial proximity with a specific region of other chromosomes.

As expected, the proximity matrices for a given chromosome in different conformational clusters vary considerably (**Fig. 5A**, second panels from the left). To quantify these differences we calculated the IPP profile for chromosome 6, defined as the total number inter-chromosomal proximities for a given chromosomal region averaged over all genome structures in a given cluster (Methods). To compare the IPP profiles between different clusters, we calculated the IppRatio, defined as the ratio of the IPP profiles in a cluster and the whole population of clustered structures (Methods). IppRatios of different clusters vary substantially (**Fig. 5A**, right profile panels). As expected, the IPPRatio of chromosome 6 in cluster 4 shows reduced exposure to inter-chromosomal interactions across the entire chromosome, due to the more compact chromosome structure observed in cluster 4 in comparison to those in other clusters (**Fig. 5A**). Indeed, cluster 4 shows the overall lowest IPPRatio values. Overall, cluster 8 shows the highest IPPRatio. However, there are differences for individual domains in each cluster. For instance, domain 3 in cluster 5 shows higher IPPratio then the corresponding region in cluster 8 (region indicated by green bar in **Fig. 5A**). Interestingly, chromosomes in different clusters favor interactions to different chromosomes (**Fig. 5B**). Chromosome 6 in cluster 8 shows the highest averaged IppRatio and thus substantially increased inter-chromosomal interactions with chromosomes 2, 21 and 8, while chromosome 6 in cluster 5 shows increased interchromosomal interactions with chromosome 20 and 16 instead (**Fig. 5B**). For instance, **Figure 5C** compares the inter-chromosomal proximity matrix between chromosome 6 and chromosome 2 for cluster 8 and cluster 4. Chromosome 6 shows increased inter-chromosomal interactions with chromosome 2 in cluster 8 in comparison to cluster 4 (**Fig. 5C**).

**Figure 5:**
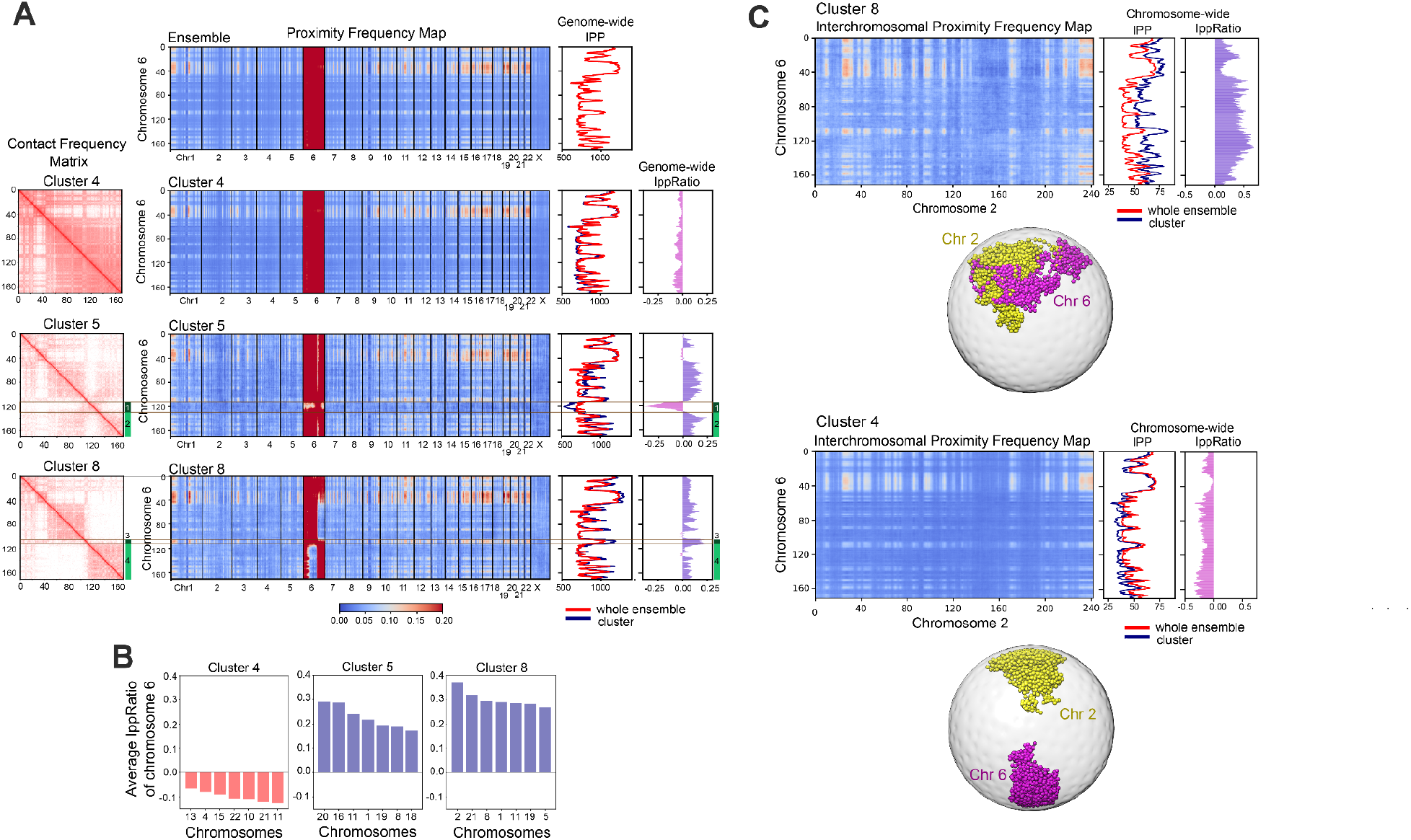
Comparison of inter-chromosomal proximity frequency map and associated features for chromosomes in different clusters. **A**, (left panels) The average contact frequency matrices of chromosome 6 calculated from all structures in different clusters. Indicated by green boxes are also 4 genomic regions, whose properties are discussed in the text. (Second panels from left) The average proximity frequency matrix between structures of chromosome 6 and structure of all other chromosomes in the genome for different clusters (Methods), (third panels from left) Inter-chromosomal proximity profile (IPP), defined as the total number of inter-chromosomal contact proximities of a genomic region with any other chromosomal region of any chromosome divided by the total number of genome structures in a cluster (Methods). The red line shows the genome-wide IPP profile calculated from the whole ensemble of structures, while the blue line shows the IPP profiles calculated from the structures in each cluster. (Forth panels from left) IppRatio, defined as the log ratio of IPP values in a cluster over the IPP value calculated from the ensemble of all clustered structures. Each row of panels shows these properties calculated from different clusters, namely clusters 4, 5 and 8. **B**, Ranking of the average IppRatios between chromosome 6 and the other chromosomes in different clusters. Only the 7 top ranked chromosomes leading to the highest averaged IPPRatio are shown. **C**, (top left panel) Interchromosomal proximity frequency map between chromosome 6 and chromosome 2 calculated from structures in cluster 8 and (bottom left panel) and structures in cluster 4. (Top middle panel) IPP profile of chromosome 6 considering interactions only to structures of chromosome 2 in cluster 8 and (bottom middle panel) in cluster 4. (Top right panel) IppRatio profiles between chromosomes 6 and 2 in cluster 8 and (Bottom right panel) in cluster 4. Also shown are representative structures of chromosome 6 and 2 in cluster 8 (top) and cluster 4 (bottom).

### Chromosome clusters can be validated by single cell Hi-C experiments

We further assessed our findings by single cell Hi-C (sci-HiC) data of GM12878 cells^10^, containing more than 11,000 single cell contact maps, each with relatively low coverage (on average 3,879 contacts per cell in 200kb resolution) (**Fig. 6AB**). To increase the relatively low contact coverage, we applied the scHiCluster method^57^, a single cell Hi-C imputation method based on linear convolution and random walk algorithms (**Fig. 6B**). About 8,500 imputed single cell contact matrices of chromosome 6 could then be classified into clusters based on their similarity to the average contact maps of our detected clusters (Methods) (**Fig. 6CD**). The average contact frequency maps for each cluster for both, the imputed and raw sci-HiC maps, show good agreement with those from our models (**Fig. 6AB, Supplementary Fig. 8B**). To ensure that the agreement is not due to overfitting, we generated a control experiment, where contact entries in each sci-HiC contact matrix were randomly rearranged while maintaining its diagonality and the number of overall contacts. Performing the same analysis with randomized sci-HiC matrices, resulted in cluster averages that did not reproduce the contact patterns in our clusters (**Supplementary Fig. 8C**).

**Figure 6:**
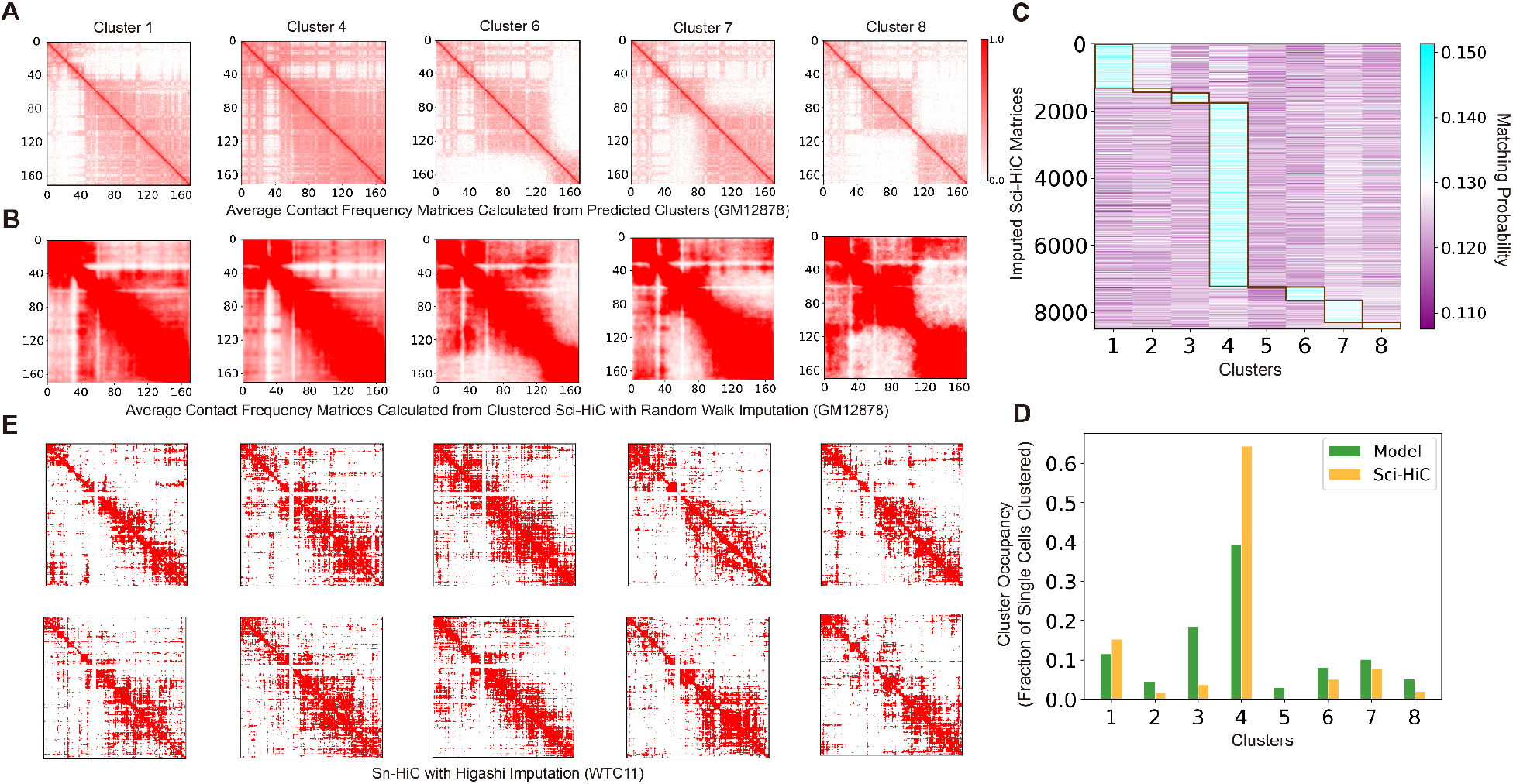
Assessment of predicted clusters by single-cell and single nucleus Hi-C data for chromosome 6. **A**, The average contact frequency matrices of clusters 1, 4, 6, 7 and 8 calculated for predicted clusters. **B**, Average contact frequency maps calculated from clustered sci-HiC contact maps^10^ imputed by convolution and random walk with restart^57^. **C**, Matching probabilities indicating the similarities of all classified sci-HiC contact matrices against all modeled clusters. **D**, Comparison of the cluster occupancy for clusters observed in our models and imputed sci-HiC data. **E**, Selected examples of sn-HiC contact matrices^36,37^ imputed by Higashi^27^ for different clusters.

Finally, we also used an available data set from single nucleus sn-HiC experiments^36,37^ of WTC11 cells (made available by the laboratory Bing Ren), which contained only 188 single cell contact maps (on average 129,499 contacts per cell in 200kb resolution) and were preprocessed by the imputation method Higashi^27^. Despite the small sample size, we found representative single cell maps of chromosome 6, with similarities in their contact patterns to those found in our clusters (**Fig. 6E**). Therefore, single cell/nucleus Hi-C data confirms the presence of predicted chromosome morphologies, even though these data do not distinguish between homologous copies. We assume that when phased single cell data becomes available the already good agreement will be further improved.

### Chromosome morphologies are conserved across cell types

Chromosome morphologies are distinguished by territory domain boundaries, which are found at a few chromosome locations and vary per cluster. To determine if chromosome morphologies and the locations of territory domain boundaries are conserved between different cell types, we applied the same clustering protocol to genome structures generated from Hi-C maps of two additional cell types, the human embryonic cell H1 hESC, and the fibroblast cell HFFc6. **Figure 7** compares the averaged contact patterns of all highly occupied conformational clusters for chromosome 6 between all the three cell types (H1 hESC, HFFc6, GM12878). In all cell types we detected very similar clusters containing chromosome structures with very similar chromosome morphologies and thus similar locations of chromosome territory domain boundaries. The only exceptions are clusters 5 and 7, which were not detected in H1 hESC cells. Thus, preferred chromosome morphologies and chromosome territory domains are largely conserved between these cell types. However, the number of structures in each cluster, i.e. the cluster occupancy, can vary between the cell types.

**Figure 7:**
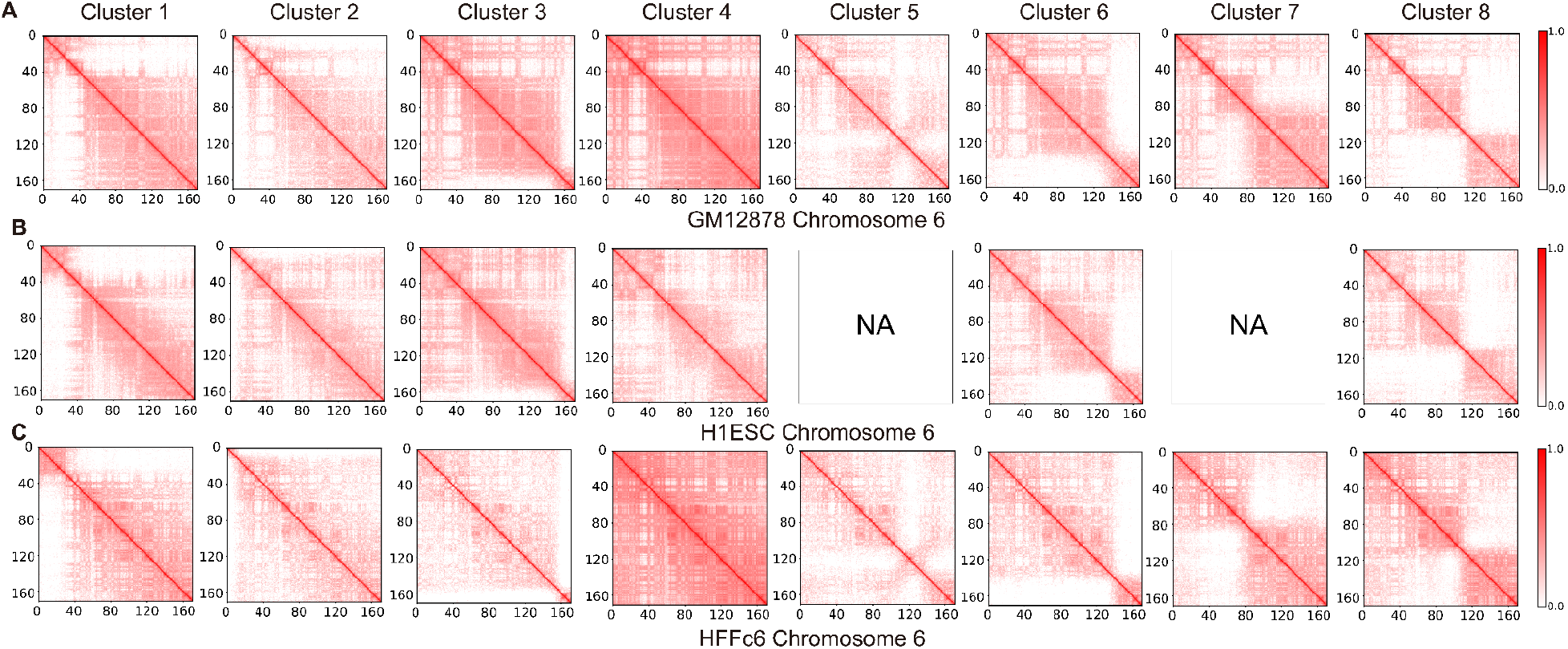
Comparative analysis of predicted clusters of chromosome 6 from genome structures of GM12878, H1-hESC and HFFc6 cells. **A**, The contact frequency matrices of the 8 predicted clusters of chromosome 6 in GM12878 cells. **B**, The contact frequency matrices of the predicted clusters of chromosome 6 in H1-hESC cells. Clusters 5 and 7 were not observed in H1-hESC cells and are indicated by “N/A”. **C**, The contact frequency matrices of the predicted clusters of chromosome 6 in HFFc6 cells.

## Discussion

Our manuscript addresses one of the key challenges in genome biology, namely, how to systematically characterize the cell-to-cell variability of whole chromosome structures and analyze the role of structural stochasticity in gene function. Due to the large dynamic variability of genome structures in a population of cells, clustering of whole chromosome and genome structures is very challenging. Traditional tools used in structural biology fail to detect functionally relevant structural similarities in chromosome morphologies, because some functionally unrelated regions can show large degree of randomness in their relative positions, overshadowing relevant structural relationships between other chromosomal regions. Subsequently, so far, little attention has been given to the heterogeneity of long-range chromosome morphologies between cells and the impact of these variations on genome function^33^. Here we present the first large-scale quantitative assessment of the 3D structural variability of whole chromosome morphologies that also considers the context of the nuclear environment. To achieve this goal, we introduce an efficient two-step clustering method that combines convolutional autoencoder with t-SNE dimension reduction to cluster large ensembles of single cell whole 3D chromosome structures, either from genome structure models or multiplex FISH imaging, into dominant conformational clusters with characteristic chromosome morphologies. Importantly, these chromosome morphologies are analyzed in their nuclear context, that is, in relation to the entire single cell diploid genome as well as relative locations to nuclear bodies and compartments. Our findings were validated by independent experiments from multiplex DNA-MERFISH imaging and single cell Hi-C.

We found that almost half of the structures for each chromosome can generally be described by 5-10 dominant chromosome morphologies, which play a fundamental role in establishing conformational variation of chromosome structures. Each morphology cluster is distinguished by a specific combination of 2 to 4 chromosome territory domains that vary between clusters. Territory domains are consecutive chromosomal regions with enhanced interactions that can be spatially separated from other territory domains. Interestingly, we found that the detected chromosome morphologies and locations of territory domain borders are largely conserved across different cell types and similar conformational clusters are found in GM12878, H1-hESC and HFFc6 cells. However, the relative cluster occupancy can vary between the cell types.

Importantly, our analysis not only discovered dominant chromosome morphologies, but also identified the relationship of chromosome morphologies with specific nuclear microenvironments of chromosomal regions. Specifically, we discovered a link between chromosome morphologies and specific subnuclear properties of chromosomal regions, such as specific preferences in nuclear positions and associations to nuclear speckles, lamina, and nucleoli. Our analysis shows that chromosome morphologies can either enhance or shield the exposure of specific chromosomal regions to the nuclear interior, exterior or nuclear speckles. Subsequently, the radial positions and speckle association frequencies for the same chromosomal region can differ substantially in different chromosome morphologies. It is known that shorter distances to nuclear speckles can enhance gene expression efficacy for some transcriptionally active genes^39–41^. Thus, by modulating the exposure of chromosomal regions to specific nuclear microenvironments, chromosome morphologies could influence chromatin function in single cells. Our work indicates that some chromosomal regions may show functional distinction in different chromosome morphologies, which could contribute to the heterogeneity of gene transcription in single cell RNA-seq experiments^58,59^. This information is crucial to uncover the role of genome structure on regulatory processes of genome function.

Most prominently we observe that some territory domains allow chromosomal regions to undergo relatively large changes in their nuclear position between different morphologies. These chromosomal regions show overall large cell-to-cell variability of their radial positions in the population of cells and are often also part of the active chromatin compartment that are embedded between extended regions of inactive B compartment chromatin. The corresponding territory domain borders are often close to transitions between the A2 and B3 Hi-C subcompartment. These regions show intermediate SON TSA-seq signals, indicative of active chromatin with a larger mean distance to nuclear speckles and lower gene expression levels than active chromatin in the A1 subcompartment, which is speckle associated in almost all cells of a population. Another interesting conclusion is that our results provide a connection between local chromatin structure properties and the presence of chromosome morphologies. For instance, we found that if a territory domain is present in a chromosome morphology, the corresponding domain boundary region shows increased chromatin fiber decompaction in comparison to the whole ensemble average. One could speculate that a decreased occupancy of cohesin (or other chromatin properties) at these locations in some single cells could favor the presence of a chromosome domain border at a specific sequence location, and subsequently could favor the presence of a specific chromosome morphology.

Finally, we also showed that genome structural models can facilitate the classification of multiplex FISH imaging data. Because of the currently relatively low coverage in multiplex FISH chromosome tracing experiments, a direct clustering of chromosome structures from experiments is challenging and can fail to detect clusters. However, as demonstrated here, it is possible to cluster chromosome structures from models generated at higher resolution. The resulting structural clusters can then facilitate the classification of chromosome morphologies in the DNA-MERFISH data. Thus, high resolution modeling of chromosome structures can also play a key role for the structural analysis of low-resolution 3D structures determined by chromosome tracing in multiplex FISH imaging experiments.

## Acknowledgements

This work was supported by the National Institutes of Health (grants U54DK107981 and UM1HG011593 to F.A). We thank Nan Hua, Wenyuan Li, Francesco Musella and Ye Wang for useful discussions. We thank Ruochi Zhang and Jian Ma for providing Higashi imputed sn-HiC of 188 cells provided by the Bing Ren lab.

## Author contributions

Y.Z. and F.A. designed research. Y.Z. performed all calculations and initial data analysis..Y.Z. and F.A. interpreted results and data analysis. Y.Z. wrote software and documentation. Y.Z., A.Y. and L.B. generated genome structures and structure features. Y.Z. and F.A. wrote the initial draft of the manuscript. Y.Z., F.A., L.B. and A.Y. edited the manuscript. All authors approved the final manuscript. F.A. secured funding.

## Methods

### Population-based modeling

We used ensemble Hi-C data with the Integrative Genome Modeling (IGM) platform^11^ to generate one 10,000 whole genome structure population for HFFc6 (raw Hi-C data from 4DN Data Portal, accession code 4DNES2R6PUEK^60^) and H1-hESC (raw Hi-C data from 4DN Data Portal, accession code 4DNES2M5JIGV^60^) cell lines, and used a previously generated and analyzed 10,000 structure population for GM12878^13^.

IGM simulates a population of structures that is compatible with the available ensemble Hi-C data by optimizing the positions of the chromatin regions. Let ***X***_*s*_ = {***x***_1*s*_, …, ***x***_*Ns*_} denote a diploid whole genome structure of *N* regions, ***x***_1*s*_ ∈ ℝ^3^ being the Cartesian coordinates of the *i*th genomic region. A population of structures is defined as a collection of *S* such structures ***X*** = {***X***_1_, …, ***X***_*S*_}. Also, let ***A*** = (*a*_*IJ*_)_*H*×*H*_ denote the Hi-C contact probability matrix, so that 0 ≤ *a*_*IJ*_ ≤1 indicates the probability that two unphased loci *I* and *J* (*I*, *J*∈{1, …, *H*}) are in contact. In the following, we will denote with (lowercase) *i* and *i*’ the two copies associated with unphased region *I* (uppercase). Our genome simulation approach numerically approximates the solution to the optimization problem 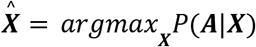, where *P*(***A***|***X***) is the probability that a population of structures ***X*** reproduces the input contact matrix ***A***. However, this poses major difficulties: first of all, it is an extremely highly dimensional maximization problem. Second, the input data *A* does not provide information on which contacts coexist within the same structure in the population and, since it is unphased, does not specify which alleles in the representation (either *i* or *i*’, *j* or *j*’) are actually in contact. In order to account for this missing information, we introduce an indicator tensor ***W*** = (*w*_*ijs*_)_*N*×*N*×*S*_, *H* ≤ *N*, such as *w*_*ijs*_ = 1(0) indicates that loci *i* and *j* are (not) in contact in diploid structure s-th. It is then essential to jointly optimize both ***X*** and ***W*** variables, i.e.

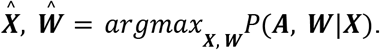

We adapted a hard Expectation-Maximization algorithm that uses a series of numeric strategies for efficient and scalable model estimation to tackle such a daunting task. We first initialize the chromatin structures in random territories, and then we start an iterative optimization, where ***W*** and ***X*** are alternatively optimized. Each iteration consists of one Assignment step (A-step), where a given subset of contacts from the input Hi-C matrix are optimally allocated across the structures (***W*** is optimized), and a Modeling Step (M-step) where the structure coordinates are optimized using Simulated Annealing Molecular Dynamics and Conjugate Gradient (***X*** is optimized). Additional batches of chromatin contacts are gradually added in each iteration, so as to improve and facilitate overall convergence. Upon convergence, a population 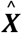 of single-cell whole genome structures are available, which are statistically consistent with the input ensemble Hi-C matrix ***A***, and also predict a number of orthogonal observables. More details on IGM formulation and implementation can be found in Boninsegna *et al*.^11^ and Tjong *et al*.^,43^.

Preliminary raw Hi-C datasets preprocessing into a 200k base pair resolution contact probability matrix was accomplished by following the protocol detailed in Yildirim *et al*.^13^.

### Genome representation

Chromosomes are represented in our models as homopolymer chains of monomers at 200-kb base-pair resolution, so that the full diploid genome is represented with N=30,332 monomers for GM12878 and N=29,838 for both H1-hESC and HFFc6. Each 200-kb chromatin region is modeled as a sphere of radius around *R*_*bead*_ = 118 *nm* in all cell lines, so that the ratio of the genome volume to the nuclear volume is 0.4^13,43^. The nuclei for GM12878 and H1-hESC are modeled as spheres of radius *R*_*nuc*_ = 5, 000 *nm*^13,43^. The nucleus for HFFc6 is modeled as an ellipsoid of semiaxes (*a*, *b*, *c*) = (7, 840 *nm*, 6, 470 *nm*, 2, 450 *nm*)^11^.

### Two-step dimension reduction

The basic aim of this study is selecting representative single cell structures from our population and studying their significance. Multiple features of a single cell structure can be extracted and calculated, such as contact matrix and distance matrix, which can be further calculated as a feature vector. Due to the high dimension of the feature vector of our 200kb model, direct classification of these feature vectors is unrealistic. We here introduce the two-step dimension reduction that preserves both efficiency and accuracy during the dimension reduction.

### Removal of unrestrained beads

For each single structure, we remove those beads that are not restrained. All beads remaining in the structure belonging to the “domain” category (not centromeres or telomeres) are considered to construct the distance matrix, while beads belonging to “cen” (centromeres or telomeres) are removed.

### Input distance matrix

The distance matrix 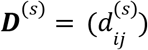 of chromosomal structure *s* is calculated by the surface-to-surface distance between bead *i* and bead *j*:

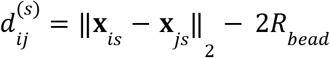

where **x**_*is*_ and **x**_*js*_ are the 3D coordinates of bead *i* and bead *j* in structure *s* and *R*_*bead*_ is the bead radius in our model. 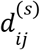 is set to be 0 if *i* = *j*. The matrix is then applied normalization to ensure the maximum entry in the matrix is 1 (dropping the structure superscript *s*):

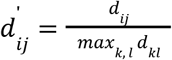

Due to the size of the layers in the convolutional autoencoder, the input matrix is then resized to multiples of 50 by bilinear interpolation so that the size of the input matrix matches the reconstructed output matrix by the autoencoder.

### Convolutional autoencoder

Convolutional autoencoders are frequently used in image classification. In this study, we regard each input distance matrix as an image, which is then regarded as the input of the input layer. The autoencoder consists of an encoder and a decoder, where the input distance matrix is the input of the encoder while the latent matrix is the output. The latent matrix is then used as the input of the decoder to generate the final output. We perform 15 epochs with batch size 200 to train the autoencoder after shuffling the input dataset. The autoencoder is implemented by the python package keras https://github.com/keras-team/keras.

#### Convolutional layer

A convolutional layer performs convolution operation to an original input over each window and constructs a new output. We use three convolutional layers in the encoder and four convolutional layers in the decoder. In the encoder, the first layer has 16 filters with a kernel with size (10, 10). The next two layers each have 8 filters with a kernel with size (10, 10) and 4 filters with a kernel with size (10, 10). We use the ReLU activation function to process the output of each convolutional layer:

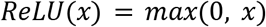

In the decoder, the first layer has 4 filters with a kernel with size (10, 10). The next two layers each have 8 filters with a kernel with size (10, 10) and 16 filters with a kernel with size (10, 10). We use the ReLU activation function to process the output of these convolutional layers. To generate the output, the last convolutional layer uses 1 filter with a kernel with size (10, 10), we use the sigmoid activation function to ensure that the values in the final output matrix are located between 0 and 1:

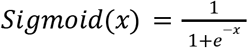

For each convolutional layer, we use the stride size (1, 1) and the same padding size to ensure the output has the same height and width as the input.

#### Max pooling layer

A max pooling layer is used to downsample an original input by calculating the maximum value in each window and generate a new value. We use three max pooling layers in the encoder. The first two layers have a pooling window with size (5, 5). The last layer has a pooling window with size (2, 2). We use the same padding size for each max pooling layer to generate the output. The stride size is the same as the window size for each layer.

#### Upsampling layer

An upsampling layer is used to up sample an original input by filling each window with the corresponding value. We use three upsampling layers in the decoder. The first layer has a sampling window with size (2, 2) and the next two layers have a sampling window with size (5, 5).

#### Latent vector

The latent vector is generated by directly flattening the latent matrix. We then use standard normalization to normalize the whole set of latent vectors to ensure that each dimension 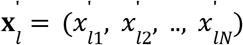 of the set of new vectors has mean 0 and standard deviation 1:

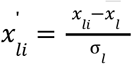

where 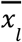 and σ_*l*_ is the mean and the standard deviation of each dimension **x**_*l*_ = (*x*_*l*1_, *x*_*l*2_, …, *x*_*lN*_) of the set of original vectors. *N* is the size of the training set.

#### Mean squared error

We use the mean squared error (MSE) to calculate the loss between the input matrix ***M***^*input*^ and the output matrix ***M***^*output*^, which is the mean Euclidean distance between the input matrix and the output matrix of the training dataset:

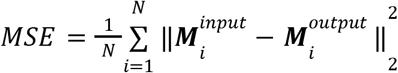

where *N* is the size of the training set.

#### Optimizer

We use an optimizer that applies the Adadelta algorithm^61^ to train the autoencoder. In comparison with other gradient descent methods, this method does not require setting of the learning rate parameter and is relatively more robust.

### T-distributed stochastic neighbor embedding

T-distributed stochastic neighbor embedding (t-SNE) is a robust and nonlinear dimension reduction method^47^. By using proper probability distributions ***P*** = (*p*_*ij*_) and ***Q*** = (*q*_*ij*_) to measure similarities between data points in both the original space and the lower dimensional space. The method facilitates the embedding by minimizing the Kullback–Leibler divergence^62^ between the two distributions by:

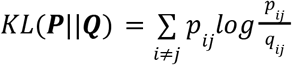

We set the dimension of the embedded space to be 2, the perplexity to be 200 and the learning rate to be 1,000.

### Principal component analysis

Principal component analysis (PCA) is a frequently used dimension reduction method which computes the principal components of the data. The method uses singular value decomposition (SVD) of the covariance matrix to construct principal components which are then used to find embedded data points in the lower dimensional space. The dimension of the embedded space is set to be 2.

### Multidimensional scaling

Classical multidimensional scaling (MDS), which is also known as Principal Coordinates Analysis (PCoA), is another nonlinear dimension reduction^48^. The classical MDS transforms pairwise distances between data points into dissimilarities and minimizes a cost function. We use Euclidean distances as dissimilarity measurement. The dimension of the embedded space is set to be 2.

### Locally linear embedding

Unlike PCA which projects data points in a linear way, locally linear embedding (LLE) is a nonlinear dimension reduction technique^49^. The method can be viewed as a collection of local PCA which preserves distances within each local neighborhood graph. The dimension of the embedded space is set to be 2.

### Isomap

Isomap, which is also a nonlinear dimension reduction method, is an extended version of MDS^50^. Specifically, the method uses geodesic distances of each local neighborhood graph as similarity measurement before performing MDS. The dimension of the embedded space is set to be 2.

### Spectral embedding

Spectral embedding (SE) is another nonlinear dimension reduction method^51^. The method uses eigenvectors of the Laplactian matrix to construct embedded data points in the lower dimensional space. The dimension of the embedded space is set to be 2. All embedding listed above including t-SNE, PCA, MDS, LLE, Isomap and SE are performed by the python package sklearn^63^.

### UMAP

Uniform Manifold Approximation and Projection (UMAP) uses knowledge of algebraic topology and simplicial complexes to perform dimension reduction^52^. The method is an increasingly frequently used nonlinear dimension reduction method which is often used to compete with t-SNE. UMAP is performed by the python package umap-learn^52^. We set the dimension of the embedded space to be 2 and the learning rate to be 1.0.

### Peak detection

In the 2D embedded conformational space, every data point represents a single structure. The structures that are closed with each other in 2D distance are more likely to have similar conformations. The next step is to sample part of the data points which are representative from the 2D distribution.

### Outlier removal

To remove outlier data points, we first calculate a pairwise distance matrix of all data points. We then generate the total distance between each data point and all the other data points by calculating the row sum of each row. The data points that have extreme total distances are removed by the 3-sigma rule. We only select data points whose row sums are within 3-sigma range 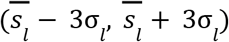, where 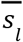 and σ_*l*_ is the mean and the standard deviation of the row sum vector *s*_*l*_.

### Kernel density estimation

Then we use bivariate kernel density estimation to calculate the probability density function of the distribution. Given a 2D independent and identically distributed sample ***X*** = (**x**_1_, **x**_2_, **x**_3_,…, **x**_*N*_), each point can be we are able to find a density function *p* so that this set of data points is sampled directly from a distribution with joint probability density function *p*:

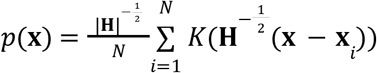

where **x**_*i*_ = (*x*_*i*1_, *x*_*i*2_)^*T*^. We choose *K* to be the gaussian kernel. The bandwidth **H** is estimated by Scott’s Rule^64^. The resulting 2D density measures how data points are distributed in the conformation space. Each local maximum of the 2D density is defined as a peak, which is a representative conformation.

### Grid approximation

We use a (1000, 1000) grid ***G*** to approximate probability density function *p*(**x**) and generate a 2D density matrix ***P***. The grid is constructed with the minimum value and the maximum value of each of the two dimensions *x*_*i*1_ and *x*_*i*2_:

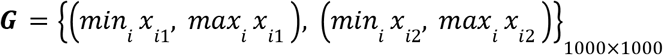

We then calculate the density value by the probability density function *p*(*x*) at each grid point and construct the density matrix ***P*** = (*p_ij_*):

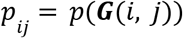

### Maximum filter

A local maximum is an entry that is larger than all its 8 neighbors in the 2D density matrix *P*. To avoid selecting multiple local maxima in a small area, however, we compare each entry with a larger range of its neighborhood. A maximum filter with size (5, 5) is applied to matrix *P* to generate another matrix 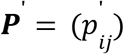. We then use exclusive disjunction (XOR) to generate matrix ***Q*** = (*q*_*ij*_) by comparing ***P*** and ***P***’ :

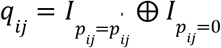

where *I*_*A*_ is the indicator function which equals 1 when *A* is true. The entries in matrix *Q* with value 1 are detected as local maxima or peaks.

### Cluster analysis

#### Boundary estimation by watershed

Considering each local maximum, we use a watershed-like approach to simulate the cluster boundary around it. We first create a density gradient with 100 levels ranging from 0 to the largest density in the 2D density matrix *P*. A set of contour lines which are polygons formed by grid points gradually change (shrink) over the density gradient. The change terminates when there is a contour in the set containing only the target local maximum, which results in our target contour line that contains only the corresponding maximum. All points surrounded by the contour line are then considered as the cluster members of the corresponding maximum. A cluster is not considered if it contains fewer than 100 members.

#### Contact frequency matrix construction

By selecting a certain number of neighbors around each peak, we are able to construct a contact frequency matrix for each peak. We estimate a path (polygon) surrounding each peak based on density. We then select all points inside the polygon as a cluster that corresponds to the peak. To calculate the contact frequency matrix ***CM***^(*a*)^ for structure *a*, we say beads *i* and *j* are in contact is structure *a* (i.e., *cm_ij_*^(*a*)^ = 1) if and only if:

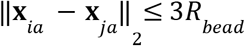

where **x**_*ia*_ and **x**_*ja*_ are the 3D coordinates of bead *i* or bead *j*. *R*_*bead*_ is the bead radius in our model. The contact frequency matrix for cluster *A*, ***CM***^(*A*)^, is calculated by the sum of all contact matrices in the cluster:

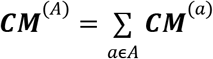

The contact frequency matrix for all structures that are classified to a cluster is calculated as:

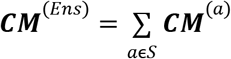

where *S* is the set of clustered structures. To enhance off-diagonal contacts, we visualize all contact frequency matrices from the models by applying transformation *log*_2_ (*cm*_*ij*_ + 1). All color bars shown together with contact frequency matrices in the figures show a ratio with regards to the maximum value.

#### Average distance matrix construction

Similarly to the construction of a contact frequency matrix, we can also construct an average distance matrix for each cluster. Each entry 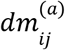 of the distance matrix ***DM***^(*a*)^ for structure *a* is calculated by the Euclidean distance between bead *i* and bead *j*:

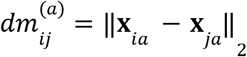

where **x**_*ia*_ and **x**_*ja*_ are the 3D coordinates of beads *i* and *j*. 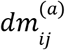 is set to be 0 if the entry is at the diagonal. After min-max normalization of each distance matrix, the average matrix for cluster *A* (which we will denote as ***DM***^(*A*)^) is calculated by the average of all matrices in the cluster:

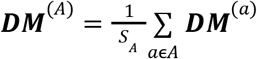

where *S*_*A*_ is the number of structures in cluster *A*. All color bars shown together with average distance matrices in the figures show a ratio with regards to the maximum value.

#### Dissimilarity measurement

##### Euclidean distance dissimilarity

We first construct two flattened distance matrices ***R***^(*a*)^ and ***R***^(*b*)^ for structure *a* and structure *b*. Each matrix contains Euclidean distances between all possible pairs of beads in each structure. The Euclidean distance between these two structures 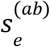 is further calculated by:

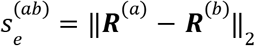

Then the final Euclidean distance dissimilarity between cluster *A* and cluster *B* is the average value of all possible pairs between these two clusters:

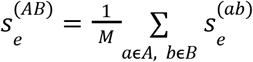

where *M* is the total number of pairs between the two clusters. To compare inter-cluster dissimilarity and intra-cluster dissimilarity, we normalize 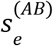 by the intra-cluster dissimilarity of cluster *A* 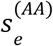:

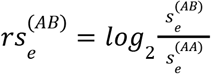

##### Gaussian dissimilarity

The calculation of Gaussian dissimilarity is adapted from Eastwood and Wolynes^54^ and Cheng et al^32^, which is an alternative way to compare pairwise distances between two structures. After generating 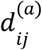 and 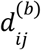 which are the Euclidean distances between bead *i* and bead *j* for both structure a and structure b, the Gaussian dissimilarity 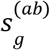 is calculated by:

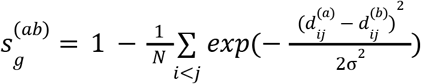

where the scaling factor σ = 8*R*_*bead*_ and *R*_*bead*_ is the bead radius in our model. *N* is the total number of pairs of beads in the structure. Similarly, the final Gaussian dissimilarity between cluster *A* and cluster *B* is the average value of all possible pairs between these two clusters:

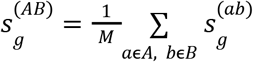

where *M* is the total number of pairs between the two clusters. To compare inter-cluster dissimilarity with intra-cluster dissimilarity, we normalize 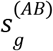 by the intra-cluster dissimilarity of cluster *A* 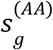:

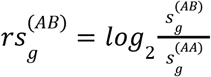

Due to computational complexity, we randomly select 200 structures from each cluster to compute the similarities above.

##### Wasserstein distance dissimilarity

We calculate both intra-cluster dissimilarity and inter-cluster dissimilarity by distance measurement to compare low intra-cluster dissimilarity with high inter-cluster dissimilarity. The Wasserstein distance *W*(*u*, *v*) measures the dissimilarity between two probability distributions *u* and *v* by:

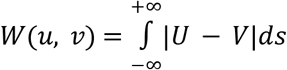

where *U* and *V* are the cumulative probability distributions of *u* and *v*^53^. To measure dissimilarity between two clusters of structures, for each pair of bead *i* and bead *j*, we obtain the 1D probability distributions for the distances between pair *i* and *j* in cluster *A* 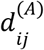 and the distances between pair *i* and *j* in cluster *B* 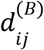, which are then used to calculate the Wasserstein distance of these two distributions. The final dissimilarity 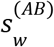 is obtained by averaging the Wasserstein distances of all possible pairs:

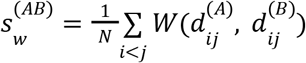

where *N* is the total number of pairs of beads in the structure. For intra-cluster dissimilarity, we randomly sample a subcluster with size 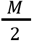 and indices *i* which is then used to calculate the final dissimilarity with its reverse subcluster with indices (*M* − *i* − 1), where *M* is the number of structures in cluster *A*. For inter-cluster dissimilarity, we directly apply the method above. To compare inter-cluster dissimilarity with intra-cluster dissimilarity, we normalize 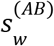 by the intra-cluster dissimilarity of cluster *A* 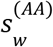:

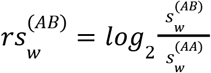

### Proximity frequency map

The calculation of a proximity frequency map is similar to the calculation of a contact frequency matrix. We select a larger range to visualize inter-chromosomal contact patterns. To calculate the proximity frequency map ***PM***^(*a*)^ for structure *a*, we define the *i*-th bead and the *j*-th bead in structure *a* forms a contact (i.e., *pm*_*ij*_^(*a*)^ = 1) if and only if:

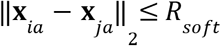

where **x**_*ia*_ or **x**_*ja*_ are the 3D coordinates of bead *i* or bead *j*. We set *R*_*soft*_ = 2, 000 *nm*. The proximity frequency map for cluster (*A*) ***PM***^(*A*)^ is calculated as:

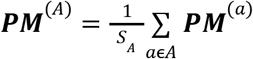

where *S*_*A*_ is the number of structures in cluster *A*. The inter-chromosomal parts of each map are shown as the average of both homologous copies, while the intra-chromosomal part is calculated from the target chromosome copy only.

### Structural features prediction

#### Radial position (RAD)

The radial position of a chromatin region *i* in structure *s* in a spherical nucleus (as GM12878) is calculated as:

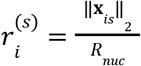

where **x**_*is*_ is the the 3D coordinates of bead *i* in structure s, and *R*_*muc*_ is the nucleus radius which is 5 μm. 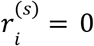 means the region *i* is at the nuclear center while 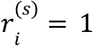 means it is located at the nuclear surface. The average radial position (RAD) of cluster *A* is the average of radial positions of all structures in this cluster:

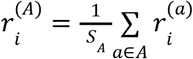

where *S*_*A*_ is the number of structures in cluster *A*. To compare against the ensemble profile, we use a log ratio comparison to show the difference. The log ratio of cluster *A* radial position against the ensemble one (RadRatio) is calculated as:

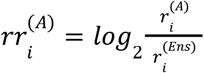

where the ensemble radial position is calculated in the same way, but for all structures that are classified to any cluster. Similarly, we can calculate all following structural features with the (*Ens*) superscript.

#### Radius of gyration (RG) (i.e., local chromatin fiber decompaction)

The local compaction of the chromatin fiber at the location of a given locus is estimated by the radius of gyration for a 1 Mb region centered at the locus. To estimate the values along an entire chromosome we use a sliding window approach over all chromatin regions in a chromosome. The radius of gyration for a 1 Mb region centered at locus *i* in structure *a*, is calculated as:

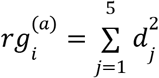

where *d*_*j*_ is the distance between the chromatin region *j* to the center of mass of the 1-Mb region. The average radius of gyration (RG) of cluster *A* is the average of radial positions of all structures in this cluster:

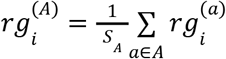

where *S*_*A*_ is the number of structures in cluster *A*. Similarly, the log ratio of cluster *A* radius of gyration against the ensemble one (RgRatio) is calculated as:

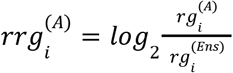

For the overall compactness of the conformation, we use all the beads to calculate the radius of gyration for structure *a*:

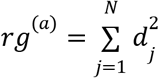

where *N* is the total number of beads in the structure, *d*_*j*_ is the distance between the chromatin region *j* to the center of mass of the whole structure.

#### Structural variability (δRAD)

The structural variability (δRAD) of region *i* in cluster *A* is calculated as:

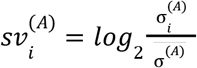

where 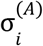 is the standard deviation of the population of radial positions of region *i* in cluster *A* and 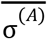 is the mean standard deviation calculated from all regions within the same chromosome of the target region. Positive values 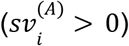 result from high cell-to-cell variability of radial position, whereas negative values 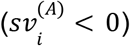 indicate low variability. The log ratio of cluster *A* structural variability against the ensemble one (δRadRatio) is calculated as:

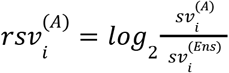

#### Building chromatin interaction networks

A chromatin interaction network (CIN) is calculated for each model and for chromatin in each subcompartment separately as follows: Each vertex represents a 200-kb chromatin region. An edge between two vertices *i*, *j* is drawn if the corresponding chromatin regions are in physical contact in the model if the spatial distance *d*_*ij*_≤4*R*_*bead*_, where *R*_*bead*_ is the bead radius in our model.

#### Identifying spatial partitions by Markov clustering

Spatial partitions of subcompartments as well as regions in A compartment with low and high structural variability are identified by applying Markov Clustering Algorithm (MCL)^65^, a graph clustering algorithm, which identifies highly connected subgraphs within a network. MCL clustering is performed for each subcompartment subgraph in each structure by using the MCL tool in the MCL-edge software^65^. The 25% smallest subgraphs (with less than 7 nodes) are discarded from further analysis to focus on highly connected subgraphs. Speckle locations are identified as the geometric center of A1 subgraphs identified by Markov clustering of A1 subgraphs. In each structure, A1 subgraphs are considered with size larger than 3 nodes.

#### Speckle distance (SpD)

The speckle distance (SpD) for region *i* is calculated by measuring the distance between the surface of each chromatin region *i* to the nearest speckle:

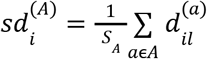

where *S*_*A*_ is the number of structures in cluster *A*, 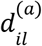 is the distance between the region *i* and the predicted nearest nuclear body location *l*. The log ratio of cluster *A* speckle distance against the ensemble one (SpdRatio) is calculated as:

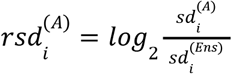

#### Speckle TSA-seq (SON TSA-seq)

Speckle TSA-seq can be viewed as an average over distances to all speckles. To predict TSA-seq signals for speckle from our models, we use the following equation:

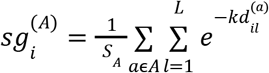

where *S*_*A*_ is the number of structures in cluster *A*, *L* is the number of predicted speckle locations in structure *a*, 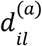 is the distance between the region *i* and the predicted nuclear body location *l*, and *k* is the estimated decay constant in the TSA-seq experiment^45^ which is set to 4 in our calculations. The normalized TSA-seq signal for region *i* then becomes:

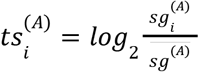

where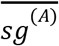is the mean signal calculated from all regions in the genome. The predicted speckles are used for distance calculations. The log ratio of cluster *A* speckle TSA-seq against the ensemble one (SON TSA-seq Ratio) is calculated as:

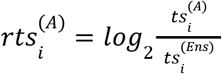

#### Speckle association frequency (SAF)

For a given 200-kb region, the association frequency to the speckle (SAF) is calculated as:

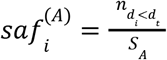

where *S*_*A*_ is the number of structures in cluster *A*, 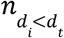 is the number of structures, in which region *i* have a distance to the speckle smaller than the association threshold *d*_*t*_. We set *d*_*t*_ to be 1,000 nm for the model and 500 nm for the DNA-MERFISH dataset^3^. For SAF calculation, we use the predicted speckle to calculate distances (see Identifying spatial partitions by Markov clustering), where we calculate distances from the surface of the region to the center-of-mass of the partition. The log ratio of cluster *A* SAF against the ensemble one (SafRatio) is calculated as:

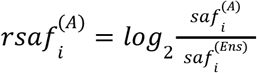

#### Inter-chromosomal proximity profile (IPP)

The calculation of inter-chromosomal proximity profile (IPP) is based on the proximity frequency map. For a given 200-kb region, the process is similar to the calculation of speckle association frequency, but we replace the distance to the smallest speckle by the contact with any inter-chromosomal regions which can be chromosome-wide or genome-wide:

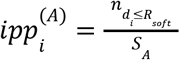

where *S*_*A*_ is the number of structures in cluster *A*, 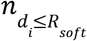 is the total number of contacts, in which region *i* is within contact range *R*_*soft*_ = 2, 000 *nm* with any target inter-chromosomal regions from the same genome structure. Every IPP is shown as the average of both homologous copies. The log ratio of cluster *A* IPP against the ensemble one (IppRatio) is calculated as:

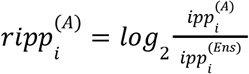

When calculating the average IppRatio of a chromosome, we calculate the mean of chromosome-wide IppRatios of the chromosome.

#### Histone modification signals and reference genes

We collected histone modification signals including H3K27ac, H3K4me1, H3K4me3, H3K9me3, H3K27me3 and H4K20me1 for GM12878 and H3K9me2 for GM23338 from the ENCODE^66^. The reference genes file for hg38 was downloaded from the UCSC Genome Browser^67^. All related signals and genes together with other structural features are shown by the Integrative Genomics Viewer (IGV)^68^.

### Cluster assessment with experimental single cell data

#### Single cell Hi-C assessment

##### Sci-HiC dataset

We collected multiple sci-HiC datasets of GM12878 from the 4DN data portal (4DNESUE2NSGS)^10^. Each dataset consists of single cell sequencing data of thousands of cells and we collected more than 11,000 single cells in total. A systematic way of massively demultiplexing single cell Hi-C is discussed in Ramani et al^10^ which applies combinatorial cellular indexing to chromosome conformation capture. We use the provided pipeline to process all collected sci-HiC datasets. Due to the large number of missing contacts, it is necessary for us to preprocess the datasets to reconstruct missing information. We adapt the preprocessing method from Zhou et al^57^. Given a raw single cell contact matrix 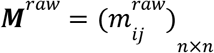, we construct a new matrix 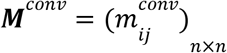 by applying convolution with filter ***F*** = (*f*_*ij*_)_(2*w*+1)×(2*w*+1)_ :

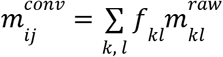

For a 200kb matrix, we set *w* = 5. In this step, we integrate the interaction information from the genomic neighbors to impute the interaction at each position. Random walk with restart is then performed to estimate contact probability between every two beads. In order to perform a random walk, a transition matrix 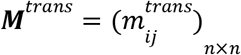 is calculated based on the contact matrix after convolution. Every entry in the original matrix is normalized by its corresponding row sum:

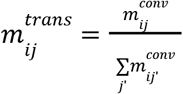

We initialize the random walk by an identity matrix ***R***_0_ so that the contact probability between every two beads is set to be 0. By applying the following recurrence formula, we are able to obtain a resulting matrix after the values converge:

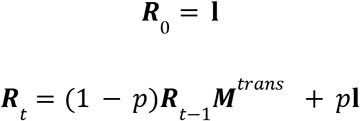

where **l** is the identity matrix and *p* is the restart probability with 0.5. We define there is a convergence when ‖***R***_*t*_ − ***R***_*t* − 1_‖_2_ ≤ 10^−6^. Each element in the resulting matrix ***R***_*t*_ after convergence indicates the probability of the random walk to reach the *j*-th node when starting from the *i*-th node. All contacts with probability larger than the 75th percentile of all probabilities in each row are chosen to convert ***R***_*t*_ into a binary matrix ***M***^*rw*^.

##### Sci-HiC assessment

For comparison of clusters, a direct way is to compare their contact frequency matrices. We define the difference matrix of cluster *A* to be:

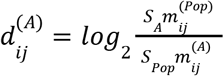

where 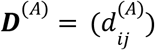 is the resulting difference matrix. 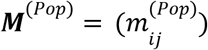 is the contact frequency matrix for the whole population calculated by contact range 2, while 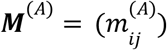 is the contact frequency matrix for the cluster. *S*_*A*_ is the cluster size while *S*_*Pop*_ is the population size. Due to the sparsity of single cell Hi-C, we preprocess each raw contact matrix by the preprocessing method above to construct a processed contact matrix. The next step is to assign each contact matrix to the clusters defined by our model. For cluster *A*, a superiority mask 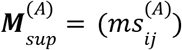 and an inferiority mask 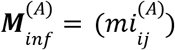 are calculated by its difference matrix ***D***^(*A*)^:

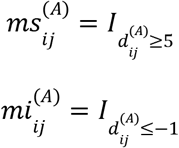

where *I_A_* is the indicator function which equals 1 when *A* is true. For each preprocessed matrix ***M***^*rw*^ from the sci-HiC population, we define the assessment score as:

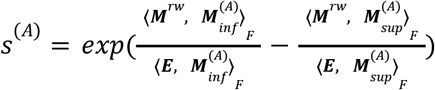

where 〈***X***, ***Y***〉_*F*_ is the Frobenius inner product between matrix ***X*** and matrix ***Y***. ***E*** is a matrix of ones. For each contact matrix, we choose the pair of masks that has the largest assessment score with the matrix and assign the matrix to the corresponding cluster. To filter matrices that are different from all clusters, we only classify matrices to a cluster *A*_1_ when 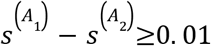, where *A*_1_ is the cluster with the largest matching score and *A*_2_ is the second largest one. The final matching probability is defined as:

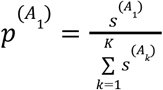

where *K* is the total number of clusters. Similarly, a contact frequency matrix can be generated using all inferred sci-HiC matrices classified to each cluster. To preserve symmetry, we symmetrize each contact frequency matrix by selecting the minimum number of contacts between pairs (*i*, *j*) and (*j*, *i*).

##### Sci-HiC control dataset

A control dataset is generated to ensure our assessment procedure is not classifying artifacts and false signals. For each sci-HiC contact matrix, we randomly rearrange all the entries while maintaining its diagonality and the total number of contacts to construct a sudo single cell contact matrix. We apply this process for every sci-HiC matrix and construct a control dataset in the end. The same assessment procedure is then applied to the control dataset.

##### Sn-HiC dataset

In total, 188 WTC-11 sn-HiC matrices were obtained from the 4DN data portal (4DNESF829JOW and 4DNESJQ4RXY5)^36,37^. Higashi^27^ is then used to impute missing contacts from the raw contact matrices. All contacts with probability larger than 0. 003*p_max_* are chosen to convert each imputed matrix into a binary matrix, where *p_max_* is the maximum probability of the imputed matrix.

#### Imaging assessment

##### DNA-MERFISH dataset

We process the DNA-MERFISH datasets from Su et al^3^ which includes high-resolution coordinates of chromosome 2 from 3,029 copies and low-resolution coordinates of chromosome 6 from 7,336 copies. For chromosome 6, we also process distances of imaged genomic regions to the nearest detected speckle. All datasets are preprocessed by linear interpolation to remove missing values if applicable. We remove copies without valid values in coordinates and in speckle distances.

##### DNA-MERFISH assessment

The preprocessed DNA-MERFISH coordinates can be then used for assessment. Each single structure is used to calculate a distance matrix ***DM*** in the same way stated above. To compare ***DM*** with the average distance matrix ***DM***^(*A*)^ for cluster *A*, we first downsample ***DM***^(*A*)^ by selecting the beads that are mapped by the DNA-MERFISH coordinates. Then we flatten both matrices by extracting the upper triangular parts and normalizing them by min-max normalization to generate two distance vectors ***R*** and ***R***^(*A*)^. We define the assessment score as:

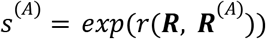

where *r*(**x**, *y*) measures the Pearson’s correlation coefficient between vector **x** and vector *y*. For each distance matrix, we choose the average distance matrix that has the largest assessment score with the matrix and assign the matrix to the corresponding cluster. To filter matrices that are different from all clusters, we only classify matrices to a cluster *A*_1_ when 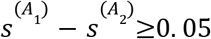, where *A*_1_ is the cluster with the largest matching score and *A*_2_ is the cluster the second largest one. The final matching probability is defined as:

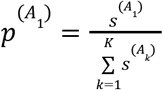

where *K* is the total number of clusters. Similarly, an average distance matrix can be generated using all DNA-MERFISH distance matrices classified to each cluster.

### Data visualization

All chromosome structures are visualized by Chimera^69^.

