## Supplementary Information for "Conformational analysis of chromosome structures reveals vital role of chromosome morphology in gene function"

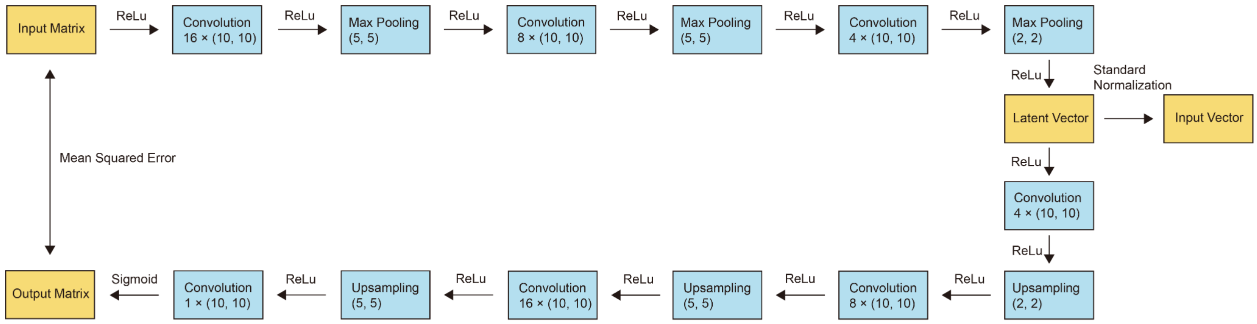


**Supplementary Figure 1: Architecture of the autoencoder** The architecture of the autoencoder consists of an encoder and a decoder. The encoder consists of three convolution layers and three max pooling layers. The decoder consists of four convolution layers and three upsampling layers. The loss between the input and the output is measured by the mean squared error.

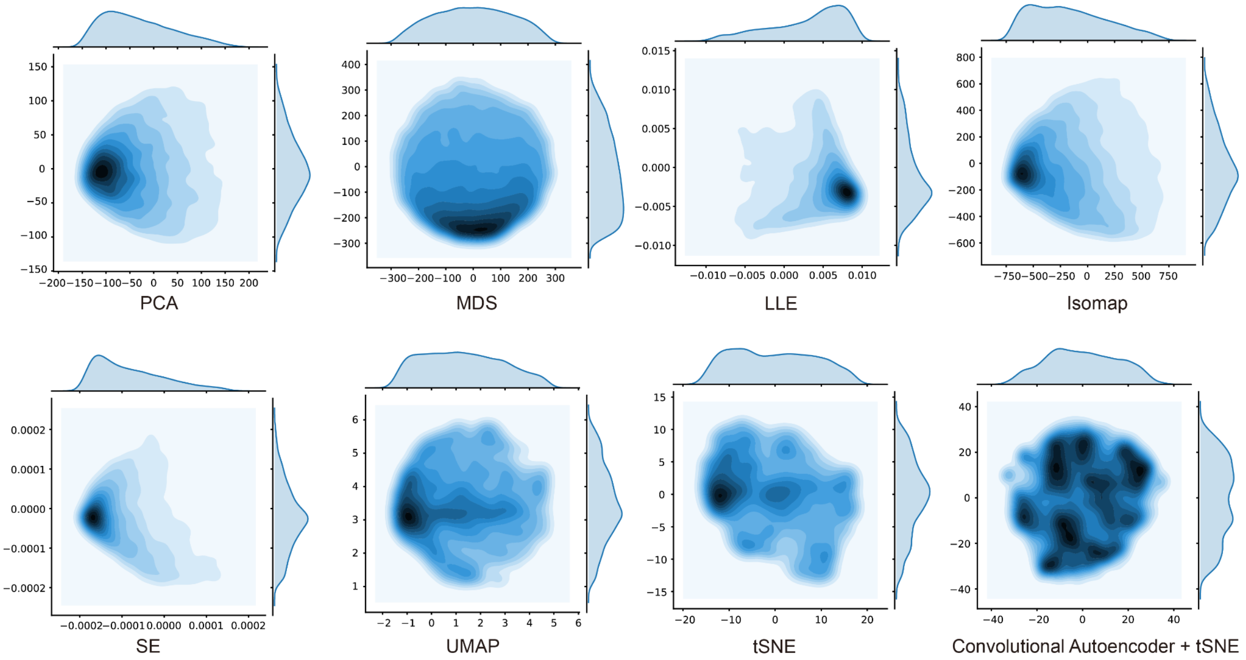


**Supplementary Figure 2: Comparison of different dimension reduction methods** Visualization of different dimension reduction methods: PCA, MDS[1], LLE[2], Isomap[3], SE[4], UMAP[5], tSNE[6]. Each method uses the same input data (the distance vectors derived from the distance matrices of GM12878 chromosome 17). After the embedded data points are obtained, we visualize the distribution by bivariate kernel density estimation. In addition we also plot the two-step dimension reduction (Convolutional Autoencoder + tSNE) proposed in this study. Note that only the two-step dimension reduction is able to generate balanced clusters.

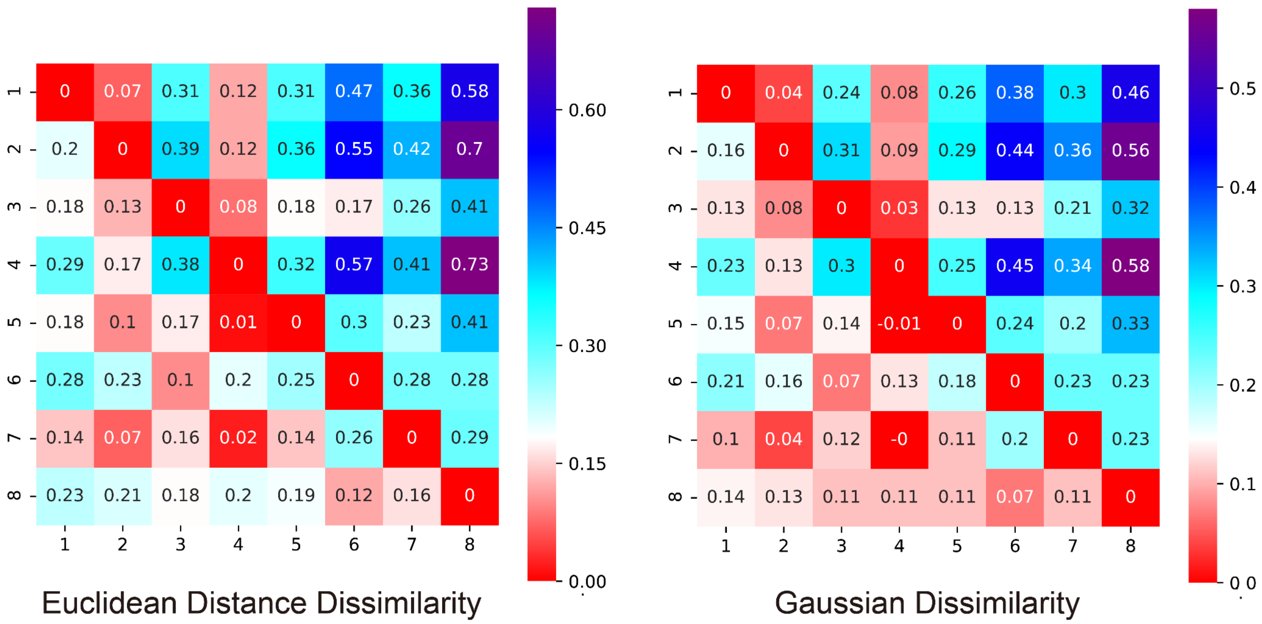


**Supplementary Figure 3: Comparison of different pairwise dissimilarity measurements on GM12878 Chr6** Pairwise dissimilarity between the 8 clusters. The dissimilarity matrices are calculated by measurements of Euclidean distance dissimilarity and Gaussian dissimilarity[7,8]. Each entry represents the log fold ratio between the inter-cluster dissimilarity and the intra-cluster dissimilarity, where positive values indicate the inter-cluster dissimilarity is larger than the intra-cluster dissimilarity.

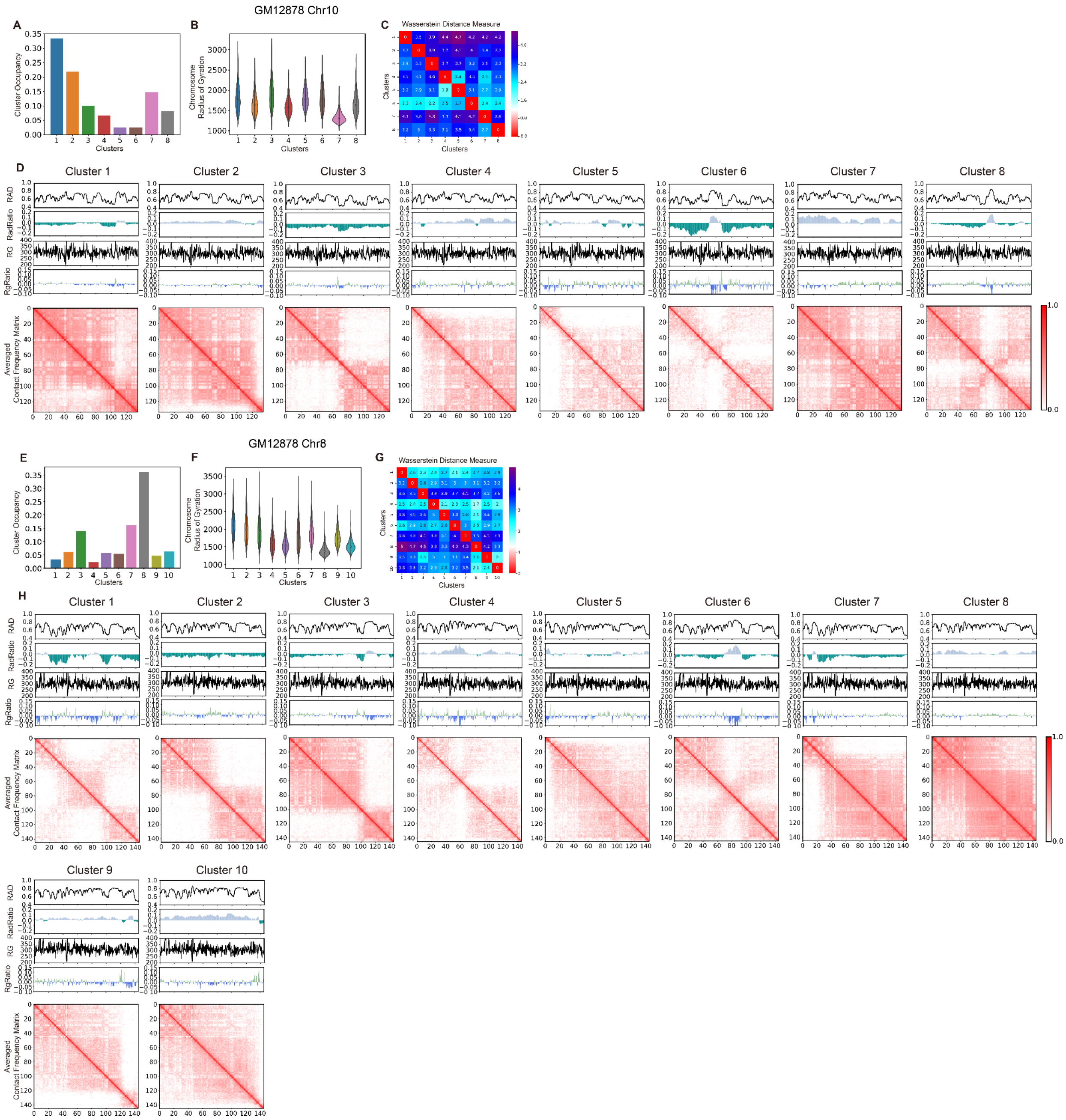


**Supplementary Figure 4: Evaluation of the method's performance on GM12878 Chr10 and Chr8 A,** The cluster occupancy of the 8 predicted clusters of Chr10. **B,** The distributions of chromosome radius of gyration for the 8 predicted clusters of Chr10. **C,** Pairwise dissimilarity between the 8 clusters of Chr10. The dissimilarity matrix is calculated by the measurement of Wasserstein distance[9]. Each entry represents the log fold ratio between the inter-cluster dissimilarity and the intra-cluster dissimilarity, where positive values indicate the inter-cluster dissimilarity is larger than the intra-cluster dissimilarity. **D,** The RAD, RadRatio, RG, RgRatio and contact frequency matrix from each of the 8 clusters of Chr10 predicted by the two-step dimension reduction method. **E,** The cluster occupancy of the 10 predicted clusters of Chr8. **F,** The distributions of chromosome radius of gyration for the 10 predicted clusters of Chr8. **G,** Pairwise dissimilarity between the 10 clusters of Chr8. The dissimilarity matrix is calculated by the measurement of Wasserstein distance. Each entry represents the log fold ratio between the inter-cluster dissimilarity and the intra-cluster dissimilarity, where positive values indicate the inter-cluster dissimilarity is larger than the intra-cluster dissimilarity. **H,** The RAD, RadRatio, RG, RgRatio and contact frequency matrix from each of the 10 clusters of Chr8 predicted by the two-step dimension reduction method.

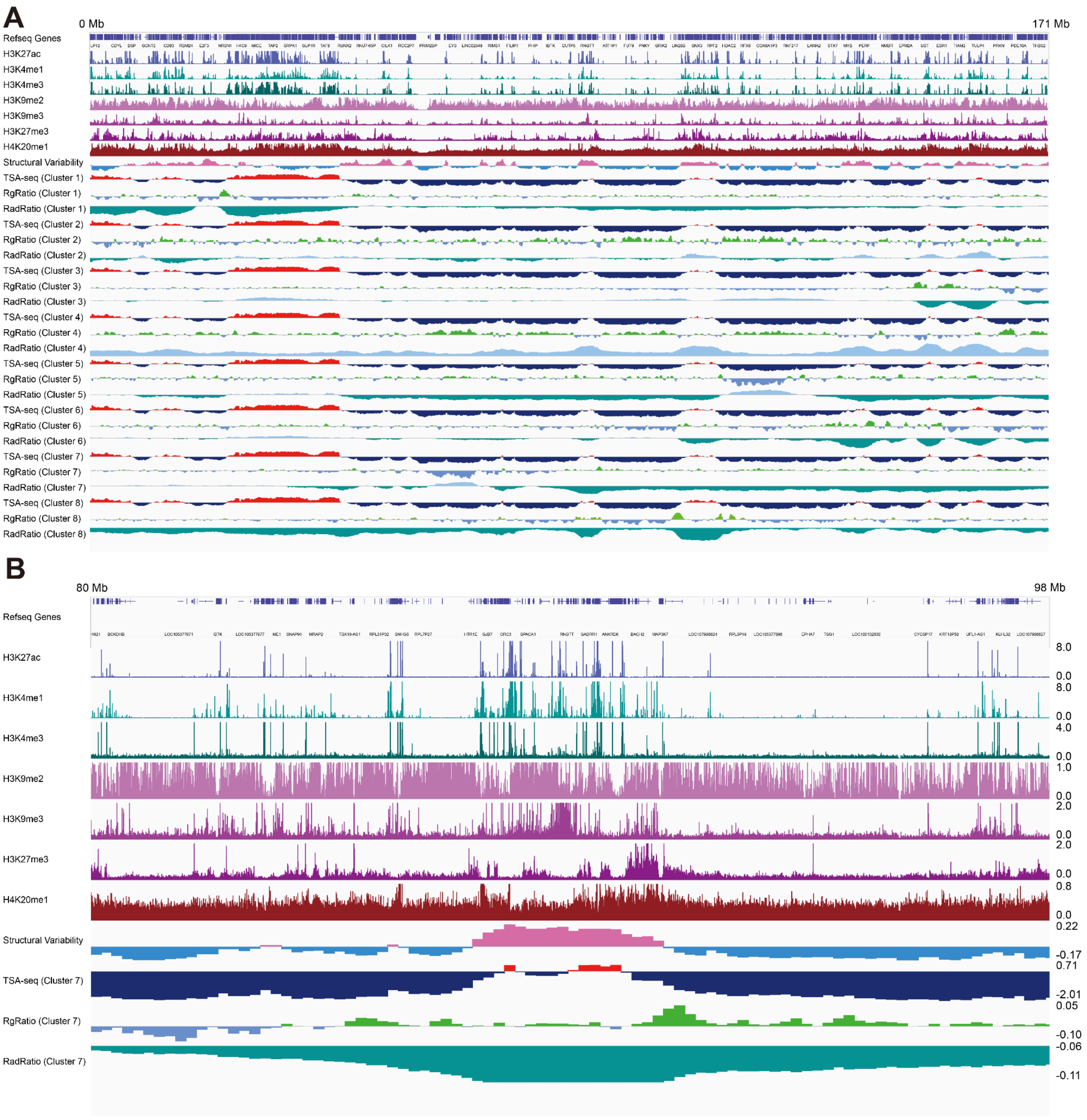


**Supplementary Figure 5: Territory domains showed together with different regulation marks on Chr6 A,** From the top to the bottom, the displayed features are refseq genes, H3K27ac, H3K4me1, H3K4me3, H3K9me2, H3K9me3, H3K27me3, H4K20me1, the ensemble structural variability, TSA-seq, RgRatio and RadRatio for all clusters. **B,** From the top to the bottom, the displayed features are refseq genes, H3K27ac, H3K4me1, H3K4me3, H3K9me2, H3K9me3, H3K27me3, H4K20me1, the ensemble structural variability, TSA-seq, RgRatio and RadRatio for cluster 7.

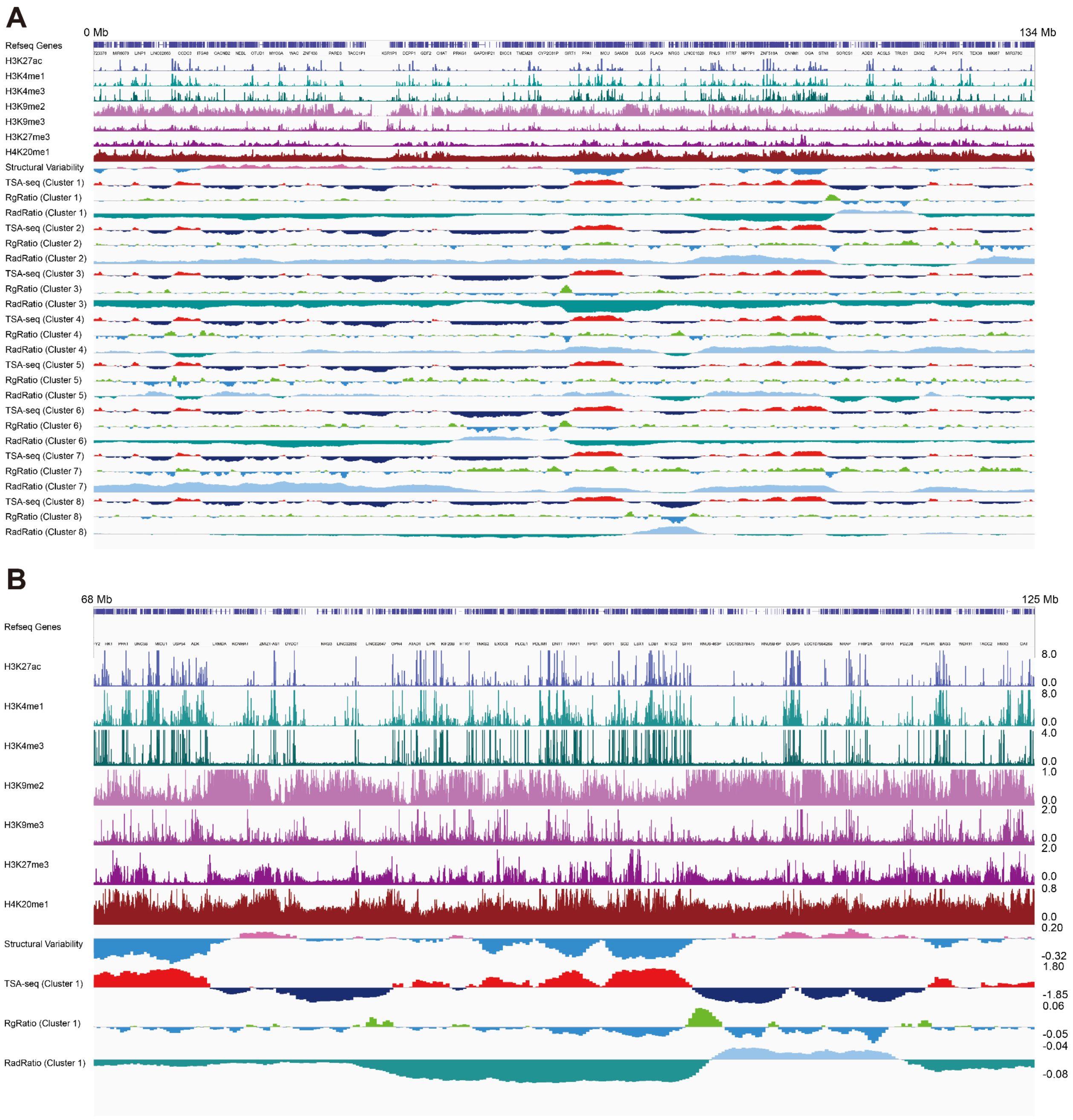


**Supplementary Figure 6: Territory domains showed together with different regulation marks on Chr10 A,** From the top to the bottom, the displayed features are refseq genes, H3K27ac, H3K4me1, H3K4me3, H3K9me2, H3K9me3, H3K27me3, H4K20me1, the ensemble structural variability, TSA-seq, RgRatio and RadRatio for all clusters. **B,** From the top to the bottom, the displayed features are refseq genes, H3K27ac, H3K4me1, H3K4me3, H3K9me2, H3K9me3, H3K27me3, H4K20me1, the ensemble structural variability, TSA-seq, RgRatio and RadRatio for cluster 1.

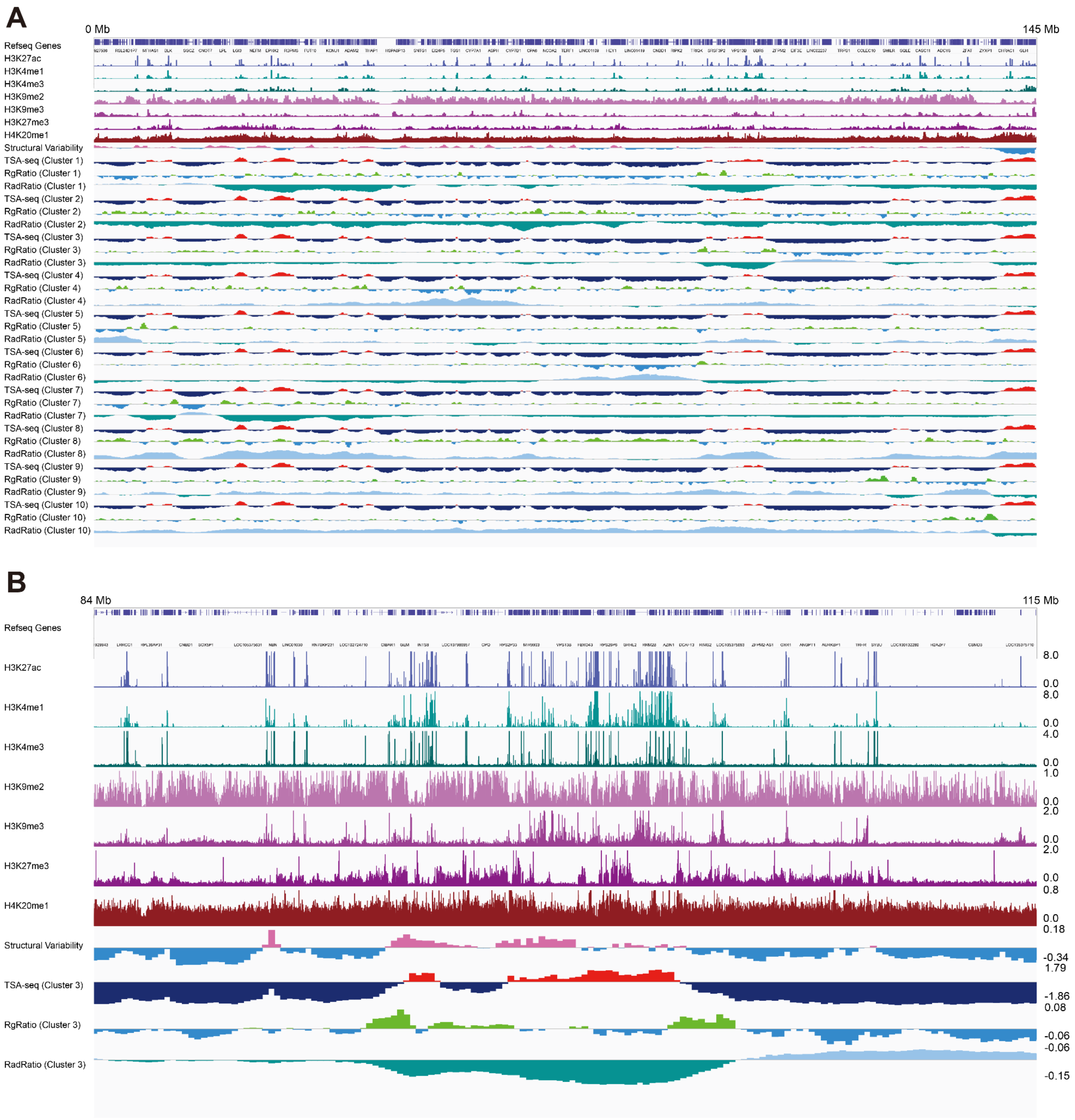


**Supplementary Figure 7: Territory domains showed together with different regulation marks on Chr8 A,** From the top to the bottom, the displayed features are refseq genes, H3K27ac, H3K4me1, H3K4me3, H3K9me2, H3K9me3, H3K27me3, H4K20me1, the ensemble structural variability, TSA-seq, RgRatio and RadRatio for all clusters. **B,** From the top to the bottom, the displayed features are refseq genes, H3K27ac, H3K4me1, H3K4me3, H3K9me2, H3K9me3, H3K27me3, H4K20me1, the ensemble structural variability, TSA-seq, RgRatio and RadRatio for cluster 3.

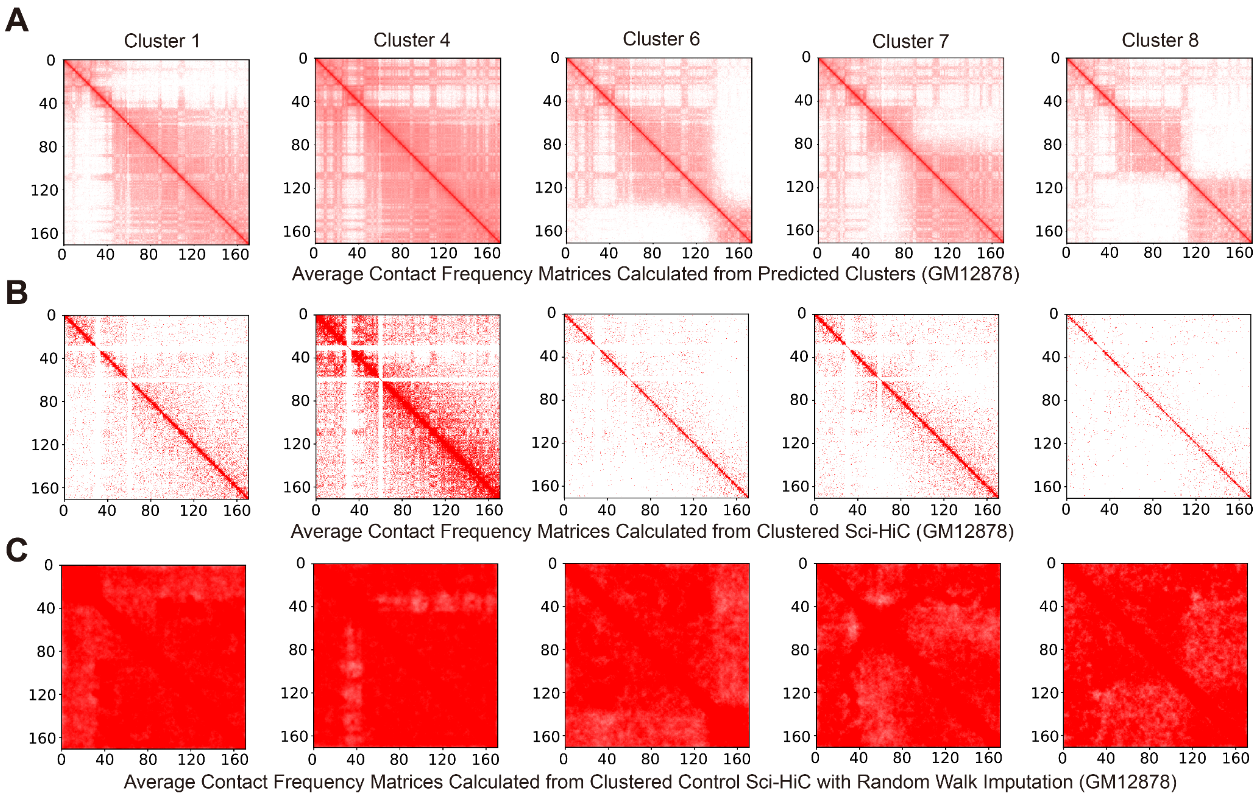


**Supplementary Figure 8: Results of the assessment of modeled clusters shown by raw and control sci-HiC data on Chr6 A,** The contact frequency matrices of clusters 1, 4, 6, 7 and 8. **B,** Results of the imputed sci-HiC assessment for different clusters. The contact frequency matrices are constructed by raw sci-HiC contact matrices[10]. **C,** Results of the imputed control sci-HiC assessment for different clusters. The contact frequency matrices are constructed by control sci-HiC contact matrices imputed by convolution and random walk with restart[11].
